# Hyperdominance and rarity in Amazonian secondary forests

**DOI:** 10.1101/2025.11.25.690408

**Authors:** Fernando Elias, Joice Ferreira, Erika Berenguer, Rodrigo Oliveira do Nascimento, Leonardo Miranda, Divino Vicente Silvério, Ima Célia Guimarães Vieira, Luiz E. O. Aragão, Gustavo Schwartz, Jos Barlow

## Abstract

Recent studies have shown that a few tree species dominate abundance and carbon storage in undisturbed Amazonian forests. However, this pattern remains poorly understood in forests regrowing after deforestation. Understanding dominance patterns in secondary forests is important as it could improve insights into their functioning and guide management and restoration efforts. We used unweighted and novel plot-weighted approaches to assess hyperdominance in abundance and carbon storage of trees and palms during secondary forest recovery, using data from 102 plots across four regions in the eastern Amazon, separated by an average distance of 600 km (range: 150-1,000 km). Secondary forests show extreme hyperdominance in all approaches, with just 16-25 species (∼3.6-5.6%) accounting for over 50% of abundance and carbon storage. A small subset of species (∼9%) were found across all regions and successional stages, highlighting their key role in ecosystem structure and function during forest regrowth. *Annona exsucca*, *Cecropia palmata*, *Tapirira guianensis*, Inga alba, *Attalea maripa*, *Amphiodon effusus*, *Banara guianensis*, and *Vismia guianensis c*onsistently emerged as hyperdominant across stem size classes, approaches, and metrics, jointly accounting for roughly 16-27% of total abundance and carbon in Amazonian secondary forests. For larger stems >10cm DBH, hyperdominant species exhibited lower wood density compared to non-hyperdominant species across all successional stages, suggesting these specialist pioneers have distinct ecological strategies from the rest of the community. Our findings indicate that secondary forest functioning can be largely predicted by a relatively limited set of species, providing a framework for future research ranging from ecophysiology to remote sensing of canopy traits and restoration planning.

## 1 INTRODUCTION

Tropical forests are at the forefront of global environmental change, with high rates of deforestation and degradation accompanied by impacts from climate change and climatic extremes (Carle et al., 2025; Lapola et al., 2023). All those impacts are leading to high levels of species turnover, with marked shifts in composition and functional traits (Pereira et al., 2025; Pinho et al., 2024). Even undisturbed tropical forests are changing as the climate warms and patterns of precipitation change (Bennett et al., 2023; Esquivel-Muelbert et al., 2019). Understanding how tropical forests are responding to global environmental change is a major research challenge given the incredible species diversity, with estimates of 10-16,000 tree species for the Amazon basin alone (ter Steege et al., 2013). However, species contribute unequally to forest structure and key ecological functions across the basin.

In the Amazon, there is a clear pattern of hyperdominance in primary forests, where just 182 of tree species account for 50% of all individuals and 227 species are responsible for 50% of aboveground carbon stocks (Fauset et al., 2015; ter Steege et al., 2013), with these patterns remaining consistent across forest strata (Draper et al., 2021). In contrast to our knowledge of hyperdominance in undisturbed primary forests, there is no assessment of hyperdominance patterns in secondary forests – i.e. those regrowing on areas that were previously deforested (Chazdon, 2014). Amazonian secondary forests are widespread, covering 180,215 km² in 2017 and storing 391.5 Tg C (Smith et al., 2021). These forests also serve as essential habitats for a wide range of fauna and flora species (Jakovac et al., 2022; Lennox et al., 2018). As such, assessing hyperdominance patterns in secondary forests is particularly relevant due to their prevalence (Nunes et al., 2020; Smith et al., 2021) and high conservation relevance (Lennox et al., 2018; Smith et al., 2023).

If there is a hyperdominance pattern in secondary forests, it is unlikely to match that observed in primary forests. First, these forests host a distinct species composition dominated by pioneers and light-dependent species adapted to environmental conditions at the initial stages of succession (Chazdon et al., 2010). A persistent selection for a small subset of pioneers may increase levels of hyperdominance compared to primary forests. Second, species turnover occurs as succession progresses in secondary forests (Poorter et al., 2023). Although pioneer species may persist over long successional periods (Fernandes Neto et al., 2019; Mesquita et al., 2001), their gradual replacement by shade-tolerant species could reduce hyperdominance across landscapes and successional stages. Third, the successional trajectories of secondary forests are influenced by a combination of environmental and anthropogenic factors, including previous land use, land occupation history, and climatic and edaphic conditions (Giles et al., 2024; Jakovac et al., 2021; Poorter et al., 2016). These factors may influence hyperdominance by promoting biotic homogenization (e.g. Solar et al., 2015) or driving divergent successional trajectories (Jakovac et al., 2021).

Beyond their ecological importance, secondary forests are also key to implementing effective large-scale ecosystem restoration aimed at mitigating climate change, halting biodiversity loss, and enhancing social well-being (Barlow et al., 2022; da Silva et al., 2023; Garrett et al., 2021). Forest restoration has a central role in Brazilian environmental legislation and various multilateral agreements that Brazil has signed. For example, under the Paris Agreement and the United Nations Decade on Ecosystem Restoration actions, Brazil has committed to restore 12 million hectares (Vieira et al., 2025). Although Brazil has yet to submit its National Biodiversity Strategies and Action Plans under the Global Biodiversity Framework, it is a signatory and has therefore committed to the restoration of 30% of degraded ecosystems by 2030 under Target 2. At regional level, significant initiatives have been proposed, such as in the state of Pará, which launched the ‘*Plano Estadual Amazônia Agora’* and the ‘*Plano de Recuperação da Vegetação Nativa do Pará*’. These initiatives outline restoration targets of up to 7.41 million hectares by 2036 to address the climate crisis and biodiversity loss (SEMAS, 2020, 2023, 2024).

Here, we draw on a dataset of 25,004 trees and palms -- including both small and large stem size classes -- sampled in 102 forest plots distributed across four regions to investigate patterns of hyperdominance in Amazonian secondary forests. Specifically, we ask:

1. Does hyperdominance occur in secondary forests, and, if so, how does it vary in terms of abundance and carbon storage when assessed using two different approaches? First, we follow the traditional approach used for Amazonian primary forests (Fauset et al., 2015; ter Steege et al., 2013) by examining the species that contribute the most, in absolute terms, and identifying those that account for 50% of the abundance or carbon storage in all secondary forest plots. However, since both abundance and carbon are strongly related to the age of secondary forests, this classical approach has some limitations when applied to secondary forests with diverse land-use histories. To address this limitation, we developed a second approach that standardizes the abundance and carbon by plot-level totals. These approaches are referred to unweighted and plot-weighted, hereafter.
2. How many of the hyperdominant species identified under the unweighted and plot-weighted approaches are also present across different geographic contexts (i.e., geographical generalists) and forest ages (successional generalists)? To explore this, we first re-classify all species based on their presence in the four regions and three age classes; then, we examine the abundance and carbon contributions of different groups of geographical and successional generalists; finally, we assess how many hyperdominants identified in question 1 are also widely distributed and successional generalists.
3. What is the taxonomic identity of hyperdominant species, and to what extent do they overlap with classifications of hyperdominance in primary forests?
4. How do hyperdominant species compare to the rest of the community in terms of their contributions to ecosystem functioning? We assess this focusing on the key trait of wood density, examining how hyperdominants and non-hyperdominants compare across the whole community and different successional time periods.

## 2 METHODS

### 2.1 Study regions

We analyze data from 102 secondary forests plots (0.25 ha each, 250 x 10 m) distributed across four macro-regions in the Eastern Amazon: Bragantina, Marabá, Paragominas and Santarém. The Bragantina region comprises the municipalities of Bragança and Capitão Poço; the Marabá region comprises the municipalities of Marabá, Canaã dos Carajás, Eldorado dos Carajás, and Parauapebas; the Paragominas region comprises only the municipality of Paragominas; and the Santarém region comprises the municipalities of Santarém, Belterra, and Mojuí dos Campos (Figure SI 1). Mean annual precipitation ranges from 1,855 mm in Marabá to 2,424 mm in the Bragantina region, with intermediate values in Paragominas (2,078 mm) and Santarém (2,182 mm). All sites have a marked dry period, which varies across regions between May and September (Figure SI 2).

### 2.2 Secondary Forest age

We determined the age of each secondary forests through interviews with landowners and/or remote sensing analyses (details in Gardner et al., 2013). Plot age at the time of the census that we use here varied between regions: Bragantina (11 to 60 years, where older plots have been extensively studied (Elias et al., 2020), Marabá (5 to 28 years), Paragominas (3 to >25 years) and Santarém (1 to >25 years).

### 2.3 Tree censuses

Tree censuses took place between 2010 and 2024. The sampling focused on two strata: larger stems with diameters at breast height (DBH) greater than 10 cm, and small stems with DBH ranging from 2 to 9.9 cm. Larger stems were measured throughout the entire 10 x 250 m plot, whereas small ones were measured in five nested 5 x 20 m subplots (Elias et al., 2022; Gardner et al., 2013). Individual identification was conducted both in the field by experienced parabotanists and by comparisons with specimens from the ‘Herbário IAN’ and ‘Herbário MG’, Brazil. The taxonomic nomenclature was reviewed and updated based on the 2020 Brazilian Flora Species List (BFG, 2018). All data used in this study are available on the ForestPlots.net platform (Lopez-Gonzalez et al., 2011).

### 2.4 Estimating abundance and carbon stocks

We calculate the aboveground biomass (AGB) of individuals using specific allometric equations for trees, palms and species of *Cecropia* genus. For trees, we applied Equation 1 (Chave et al., 2014), and for palms, we applied Equation 2 (Goodman et al., 2013). For *Cecropia* species, we used Equation 3 (Nelson et al., 1999), due the hollow stem structure that make general allometric models unsuitable.

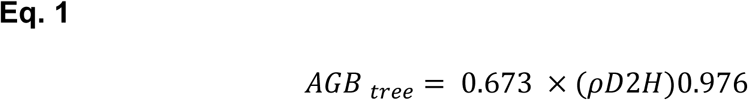

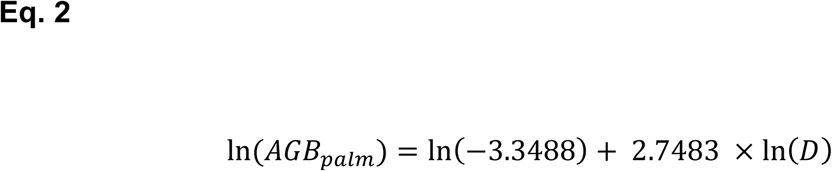

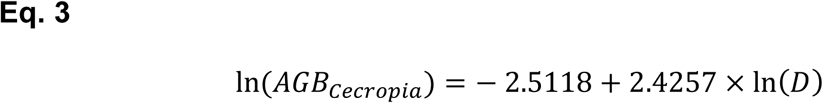

where *ρ* is wood density extracted from the Global Wood Density Database (Zanne et al., 2009); D is diameter at breast height (cm); and H is total height (m) estimated by height-diameter model (see parameters of HD model in the Table SI 1).

We estimate the carbon content using a ratio of 0.5 of AGB values (Ngo et al., 2013). We calculate plot-level carbon storage as the sum of the carbon stock of all individuals in a plot, extrapolated by hectare. Abundance was defined as the total number of individuals of each species per plot.

### 2.5 Hyperdominance approaches

To assess patterns of hyperdominance in abundance and carbon storage among our secondary forest plots, we applied two approaches, hereafter referred to as *unweighted* and *plot-weighted hyperdominance*. For the unweighted approach, we used Equation 4:

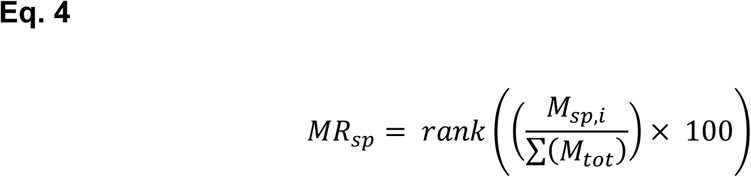

where 𝑀𝑅_𝑠𝑝_is the relative metric value for abundance (*A*) or carbon storage (*C*) for each species, ordered from the largest to the smallest; 𝑀_𝑠𝑝,𝑖_ is the metric value (*A* or *C*) for each species *i*, summed across all sampled plots; and ∑(𝑀_𝑡𝑜𝑡_) is the sum of values for each metric for the entire community across all sampled plots. Species are considered hyperdominant if their cumulative share of 𝑀𝑅_𝑠𝑝_ values accounts for at least 50% of the total community metric, following the approach commonly applied to primary forests at the biome level (cf. Fauset et al., 2015; ter Steege et al., 2013).

In the plot-weighted approach, we recognize that environmental factors (e.g., forest age, land-use history, climate, and soil characteristics) are key determinants of carbon storage and species abundance during forest recovery (Elias et al., 2022; Giles et al., 2024; Jakovac et al., 2021; Poorter et al., 2016). Unweighted approaches therefore tend to give highest values to species present in older forests or in sites with favorable land-use histories or climate conditions. To address this, we also calculated *plot-weighted hyperdominance* for both abundance (*A*) and aboveground carbon (*C*) metrics using Equation 5:

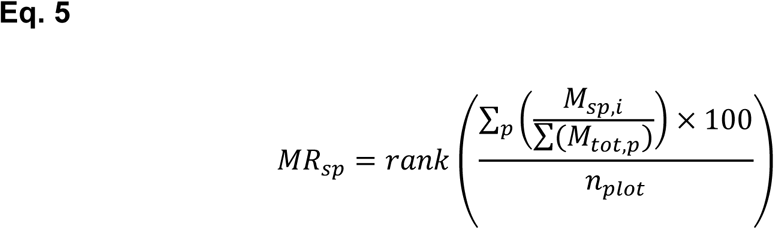

where 𝑀𝑅_𝑠𝑝_ is the plot-weighted relative metric (e.g., relative abundance or relative carbon) for each species, ordered from the largest to the smallest; 𝑀_𝑠𝑝𝑖_ is the metric value (*A* or *C*) for each species within each plot; ∑(𝑀_𝑡𝑜𝑡,𝑝_) is the total value of the metric for the community within each plot (*p*); and 𝑛_𝑝𝑙𝑜𝑡_ is the total number of sampled plots (n = 102). Hyperdominant species were defined as those ranked up to the threshold where their cumulative share of 𝑀𝑅_𝑠𝑝_ values accounts for at least 50% of the total community metric. We repeated these analyses for both approaches separately for large and small stems.

### 2.6 Statistical analysis

To compare abundance and carbon hyperdominance between the two approaches for larger and small stems, we used the relative contribution of each species across all secondary forest plots and constructed a biplot of species rank against cumulative proportion. Pearson’s correlations were used to assess the degree of convergence between abundance and carbon hyperdominance patterns across species.

To evaluate whether hyperdominant species, as defined by each approach, differ from non-hyperdominant species in their ability to occupy different successional and geographic contexts, we evaluated species occupancy across plots, stratified by the four study regions and three forest age classes (0–10, 11–20, and >20 years). We then computed the relative contribution of each species to total carbon stocks and abundance per plot and summed these values within species groups. This calculation was also performed for species restricted to a single region and age class, as well as for all possible region–age class combinations, to examine variation in species performance across space and successional gradients. We used this to further restrict the definition of hyperdominance to isolate the species present in all regions and successional stages and that jointly accounted for 50% of the total relative abundance and carbon. The variation was visualized using a heatmap, accounting for differences in occupancy among regions and age classes. Based on the premise that true secondary forest hyperdominants should be widespread across both geographic and successional gradients, we also constructed a biplot to highlight species classified as hyperdominant under both approaches, and to identify which of these occurred across all regions and successional classes.

To assess the extent to which hyperdominance classifications reflect functional variation, we analyzed and compared wood density among hyperdominant and non-hyperdominant species, across both hyperdominance approaches and successional stages. We then compared group means and confidence intervals graphically.

All analyses were conducted both on the full dataset (all individuals combined) and separately for large and small stems. Individuals not identified at the species level were excluded. All statistical analyses and graphs were performed using R software 4.5.0 at a 5% significance level (R Core Team, 2025).

## 3 RESULTS

Our assessment measured and identified 9,283 larger stems ≥10cm DBH and 15,711 small stems <10cm DBH and recorded 441 and 592 species, respectively. The total sample recorded 75 families, 285 genus and 683 species.

### 3.1 Assessing hyperdominance in secondary forests

Using the unweighted approach, just 25 species (5.6%) accounted for more than 50% of the secondary forest abundance, while 16 species (3.6%) accounted for 50% of carbon storage (Figure 1; Table SI 2). These numbers and percentages were similar using the plot-weighted approach, with 18 and 24 species (5.4 or 4%) accounting for more than 50% of the secondary forest abundance or carbon storage, respectively (Figure 1; Table SI 2). Abundance and carbon hyperdominance exhibited approximately 90% convergence, consistently maintained across both approaches and stem size classes (Figure 2, Figures SI 3).

**Figure 1.**
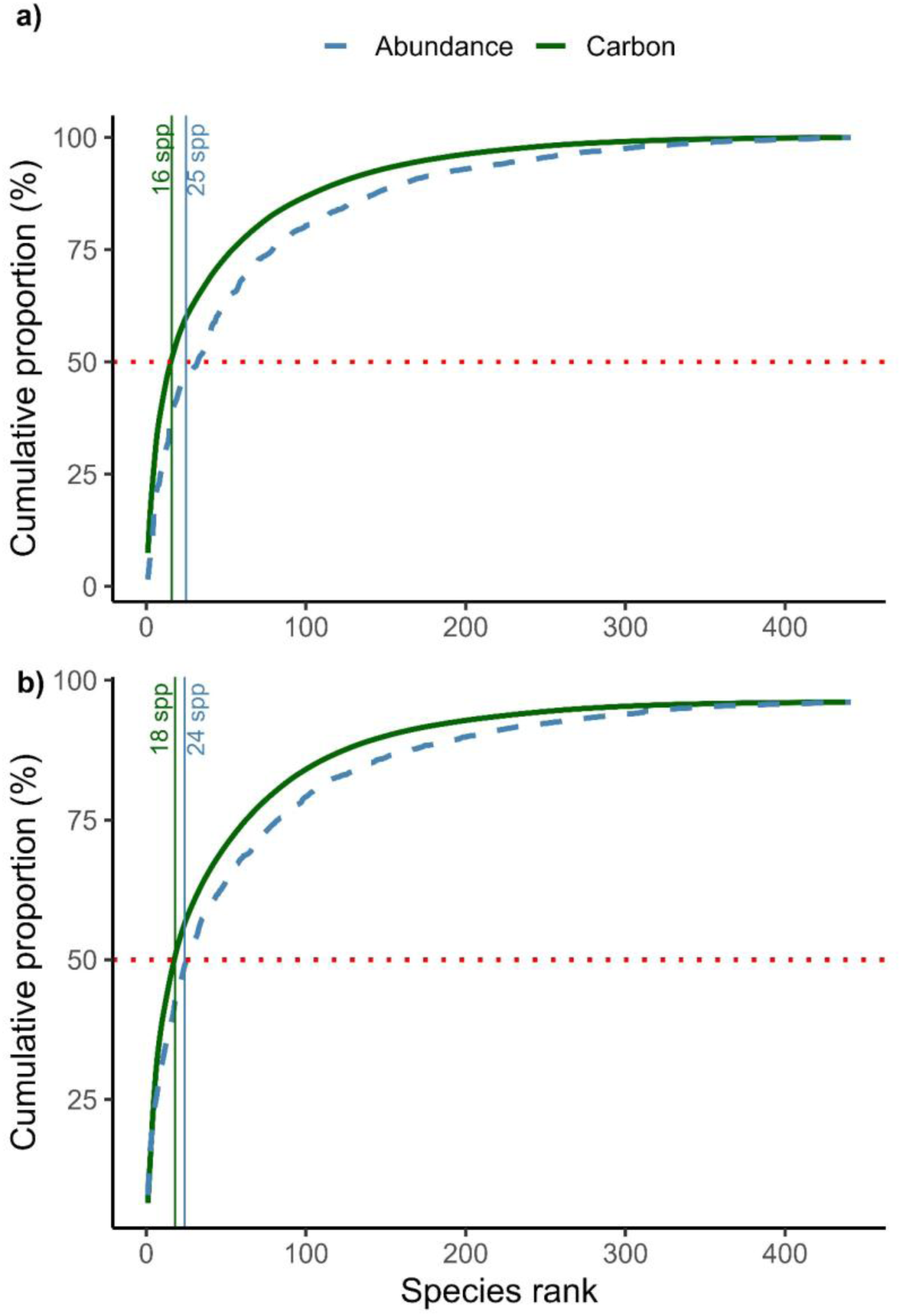
Rank of species dominating carbon storage and abundance of larger stems. Panel A shows unweighted hyperdominance based on cumulative values across all plots; Panel B shows plot-level weighted hyperdominance. The red dashed line marks the species contributing to 50% of cumulative dominance. Vertical lines indicate hyperdominant species identified by each approach and parameter.

**Figure 2.**
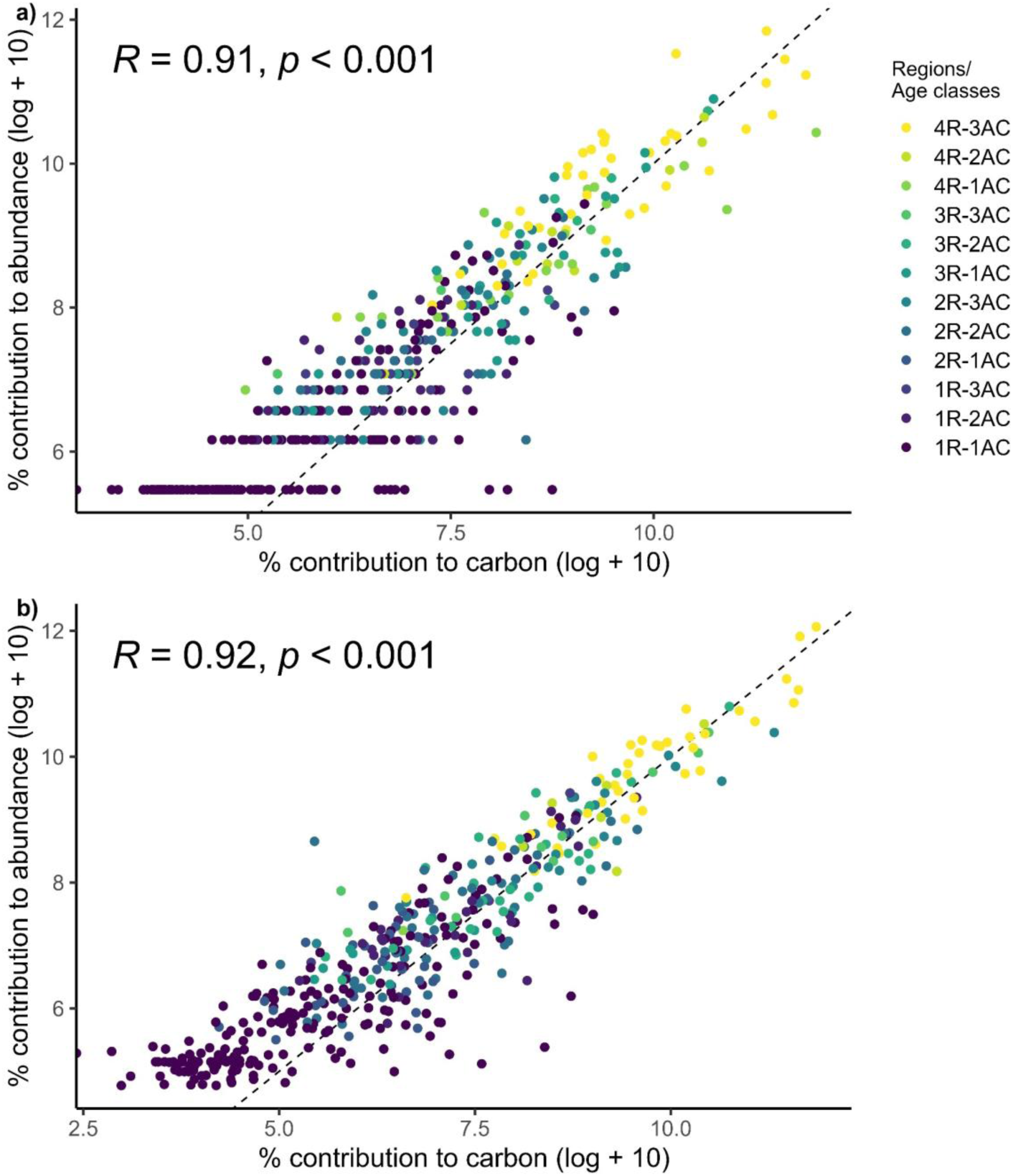
Convergence between the relative contributions to abundance and carbon for large stems in Amazonian secondary forests. Values are based on (a) unweighted and (b) plot-weighted approaches across four evaluated regions (R) and three forest age classes (AC). Both axes show log-transformed values (log + 10). The black line represents the fitted relationship trend.

### 3.2. Hyperdominant persistence across geographic contexts

Most sampled species (46.5%) are represented in single age classes and regions, while just ∼9% occurs across all regions and ages (39 spp.) among both larger and small stems. However, these species accounted for 54.4% of abundance and 54.7% of carbon storage in larger stems (Figure 3), with a slight decrease observed in smaller stems (Figure SI 5). Among hyperdominant species identified using both unweighted and plot-weighted approaches, 65.5% of them occurred across all regions and age classes in larger stems, and 28.8% in smaller stems (Figure 4, Figure SI 6).

**Figure 3.**
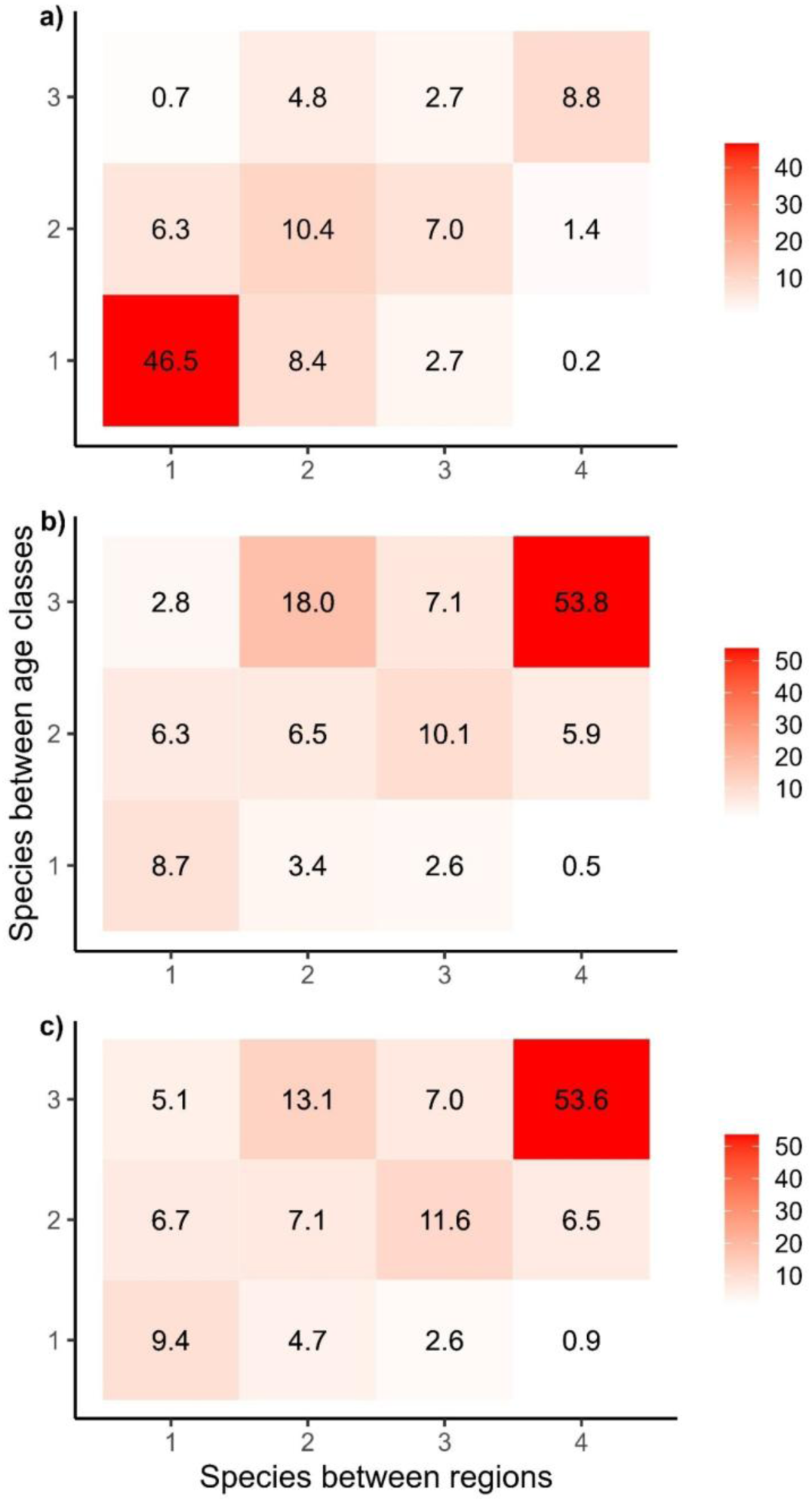
Hyperdominance among larger stems of woody species in Amazonian secondary forests. Panel (a) shows the proportion of species shared among the four evaluated regions and three forest age classes. Panels (b) and (c) show the relative contributions of carbon and abundance, respectively, for the species present in each interaction shown in panel (a).

**Figure 4.**
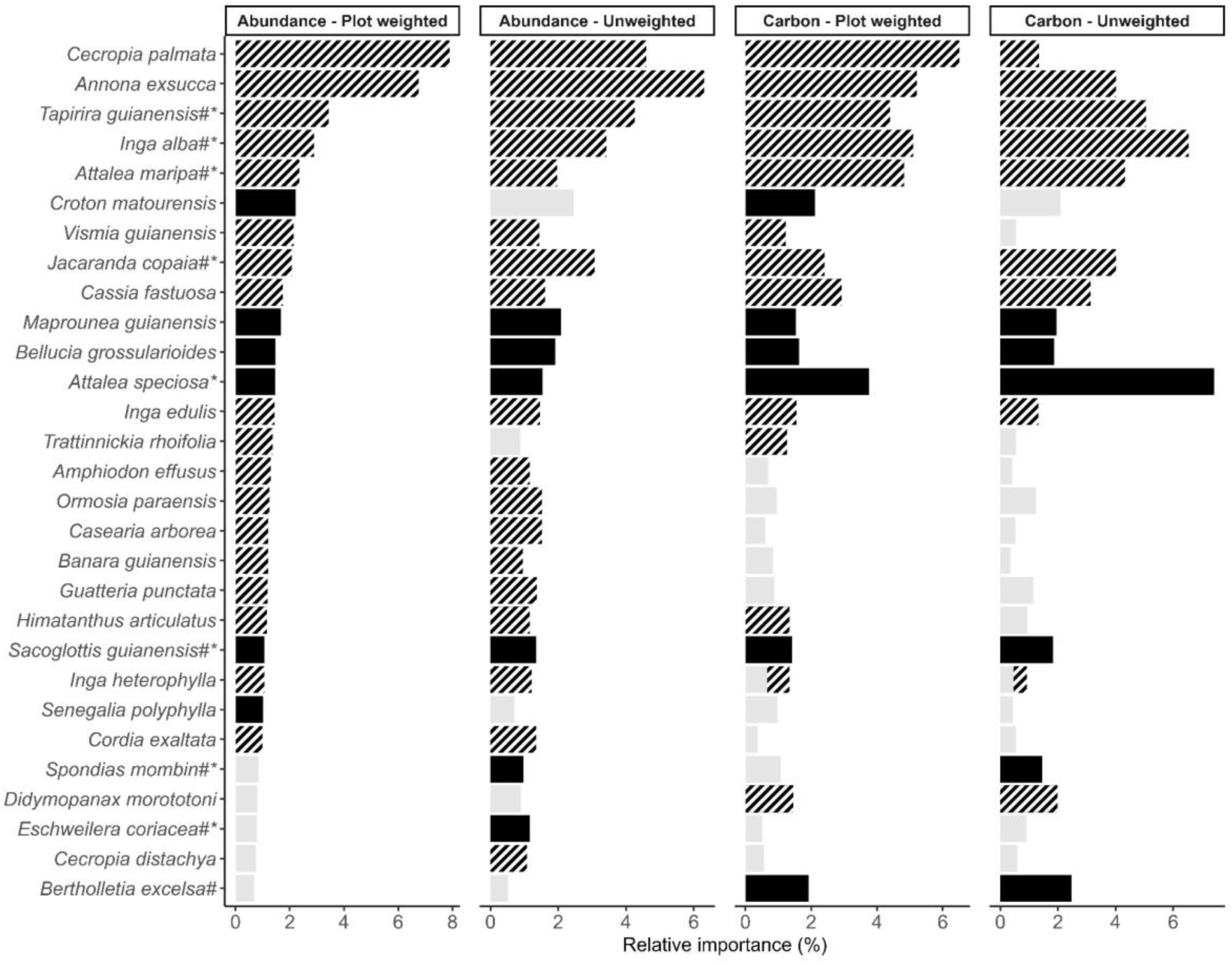
Relative contribution of hyperdominant species to carbon storage and abundance based on unweighted and plot-weighted approaches for larger stems. Black bars indicate that are hyperdominant species for carbon storage and abundance under both approaches. Black striped bars indicate species present in all four regions and age classes. Light grey bars correspond to non-hyperdominant species. Species are ordered by plot-weighted abundance. * = species classified as abundance hyperdominant by ter Steege et al. (2013); and # = species classified as carbon hyperdominant by Fauset et al. (2015).

### 3.3 Which species are hyperdominants?

The top five species classified as hyperdominant in abundance among larger stems under the unweighted approach were *Annona exsucca*, *Cecropia palmata*, *Tapirira guianensis*, *Inga alba*, and *Jacaranda copaia*, accounting for 21.6% of total abundance. Under the plot-weighted approach, four of these species (*A. exsucca*, *C. palmata*, *T. guianensis*, and *I. alba*) remained, with *Attalea maripa* replacing *J. copaia*, summing to 23.3% of total abundance (Figure 4; Table SI 2). Among smaller stems, the unweighted approach identified *Amphiodon effusus*, *Vismia guianensis*, *Banara guianensis*, *Vismia baccifera*, and *A. exsucca* as the top five hyperdominant species, accounting for 19.3% of total abundance. In contrast, the plot-weighted approach identified the same species set, except that *A. exsucca* was replaced by *C. palmata*, contributing 19.4% to total abundance (Figure SI 6; Table SI 2).

In terms of carbon, differences in species composition were observed between approaches for large stems. The unweighted approach identified *Attalea speciosa*, *I. alba*, *T. guianensis*, *A. maripa*, and *A. exsucca* as the top five hyperdominants, representing 27.3% of total carbon. The plot-weighted approach included *C. palmata*, *A. exsucca*, *I. alba*, *A. maripa*, and *T. guianensis*, with a slightly lower cumulative contribution (26%). For small stems, the unweighted approach indicated *A. effusus*, *A. exsucca*, *B. guianensis*, *V. guianensis*, and *C. palmata* as the dominant species in terms of carbon, together accounting for 16.4%. The plot-weighted approach identified the same species, though in a different order, with a combined contribution of 19.2% (Figure 4, Figure SI 6; Table SI 2).

The composition of hyperdominant species among larger and small stems varied slightly between the two hyperdominance approaches (Figure 4). For larger stems, three species (*Himatanthus articulatus*, *Trattinnickia rhoifolia*, and *V. guianensis*) classified as carbon hyperdominants under the plot-weighted approach, and two species classified as abundance hyperdominants (*T. rhoifolia* and *Senegalia polyphylla*) were not identified as hyperdominant under the unweighted approach. Similarly, although *Eschweilera coriacea*, *Cecropia distachya*, and *Spondias mombin* were classified as hyperdominant under the unweighted approach, they were not identified as abundance hyperdominants under the plot-weighted approach. Additionally, *S. mombin* was also not classified as a carbon hyperdominant under the plot-weighted approach (Figure 4).

For small stems, *Bauhinia acreana*, *Sagotia racemosa*, and *Solanum crinitum* identified as carbon hyperdominants and *Ambelania acida*, *Lindackeria paraensis*, *Miconia minutiflora*, *S. racemosa*, and *S. crinitum* identified as abundance hyperdominants were not retained under the unweighted approach. Conversely, *Zanthoxylum rhoifolium* (carbon) and *Myrcia cuprea* (abundance), classified as hyperdominant under the classical approach, were not retained under the plot-weighted approach (Figure 4, Figure SI 6; Table SI 2). Among species classified as hyperdominant for larger stems under the unweighted approach, nine carbon hyperdominants and 18 abundance hyperdominants occurred across all regions and age classes evaluated. For small stems, only 12 carbon hyperdominants and 18 abundance hyperdominants were present across all regions and age classes (Figure 4; Table SI 2).

Palms were also key contributors to carbon storage, particularly among larger stems. *A. speciosa* and *A. maripa* were the most dominant in terms of carbon storage, especially in Marabá and Santarém, whereas *A. gynacanthum* was highly abundant among smaller stems in Santarém (Figure 4, Figure SI 6; Table SI 2).

In our secondary forest plots, 14 species classified as hyperdominant for abundance and another 14 for carbon were also hyperdominant among larger stems in primary forests, including *T. guianensis*, *I. alba*, *J. copaia*, *A. maripa*, *Sacoglottis guianensis*, *S. mombin*, *Bertholletia excelsa*, and *E. coriacea*. In contrast, 13 species classified as hyperdominant among smaller stems overlapped with primary forest hyperdominants for abundance -- such as *Cupania scrobiculata*, *E. coriacea*, *T. guianensis*, and *Inga capitata* -- whereas only six species overlapped for carbon, notably *E. coriacea*, *T. guianensis*, and *Couratari stellata* (Table SI 2).

### 3.4. Hyperdominant vs. non-hyperdominant wood density

Hyperdominant species in our secondary forest plots exhibited lower wood density compared to non-hyperdominant species among larger stems (Figure 5a). In contrast, no significant differences were observed for small stems (Figure SI 7a). Wood density patterns varied between small and larger stems and between hyperdominant and non-hyperdominant species across forest age classes. Among larger stems, hyperdominant species showed no variation in wood density across age classes, in contrast to non-hyperdominant species, which exhibited lower wood density in forests up to 10 years old and higher, more stable values in forests older than 11 years. For small stems, both hyperdominant and non-hyperdominant species showed similar patterns, with younger forests (≤10 years) presenting lower wood density compared to older age classes (>11 years) (Figure 5b, Figure SI 7b).

**Figure 5.**
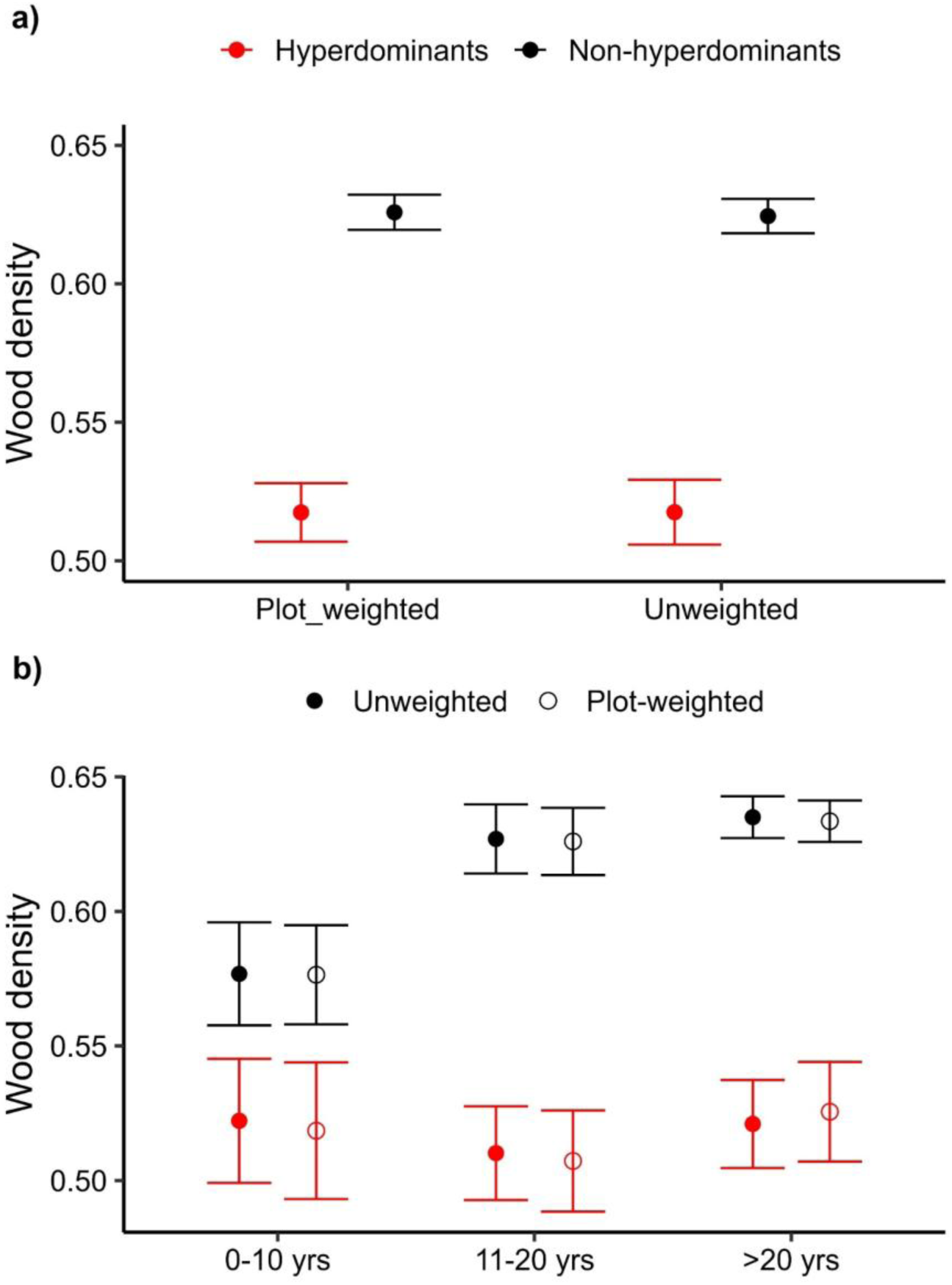
Variation in wood density among hyperdominant and non-hyperdominant species among larger stems. Panel a shows differences between the two groups across both hyperdominance approaches, while panel b shows their variation across forest age classes and both approaches. Panel (a) shows the differences between the two groups across both approaches, while panel (b) shows their variation across forest age classes under both approaches.

## 4 DISCUSSION

### 4.1 Hyperdominance in secondary forests

Our study provides the first multi-region assessment of hyperdominance in carbon storage and species abundance within tropical secondary forests. Irrespective of the hyperdominance approach, just 3 to 5.6% of the secondary forest species accounted for over 50% of the carbon storage or abundance in the plots where they occurred. Levels of hyperdominance in secondary forests are therefore similar to those observed in undisturbed Amazonian primary forests, where 1.4% of all species (227 spp.) account for total abundance (ter Steege et al., 2013), and ∼5.3% (182 spp.) store over 50% of aboveground carbon storage across the Amazon (Fauset et al., 2015).

Although our assessments are more restricted to the eastern Amazon than the pan-Amazonian assessments of primary forests, secondary forests are not uniformly distributed and are more prevalent in the ‘Arc of Deforestation’. Pará state alone has over 260,000 km² of deforested land, of which approximately 22.4% is covered by secondary forests (da Silva et al., 2023; Smith et al., 2021). Furthermore, there are some reasons to believe our patterns may extend to other Amazonian regions. First, our hyperdominant secondary forests species are widely distributed in the Pan-Amazon region. Eight species (1.8%), such as *Eschweilera coriacea* and *Inga alba*, are among the 227 species classified as hyperdominants in the Pan-Amazon (ter Steege et al., 2013). Second, five of them (e.g., *Tapirira guianensis*, *Spondias mombim*, *Didymopanax morototoni*) are among the main species with wide occurrence in successional forests, occurring in five or more floristic groups in the Neotropical region (Jakovac et al., 2022). Finally, our hyperdominant species, such as *Cecropia palmata* and *Vismia guianensis*, are also reported as important in other studies conducted in the Amazon biome, including areas near Tefé and Manaus in the Central Amazon (Fernandes Neto et al., 2019; Mesquita et al., 2001; Wieland et al., 2011).

These and other hyperdominant species in secondary forests can be considered both successional specialists and geographical generalists, as they dominate despite the wide range of forest ages, land-use intensities, and other drivers of succession represented in our study regions. This persistent hyperdominance occurs even under distinct ecological processes and regional environmental conditions that typically shape recovery trajectories (Giles et al., 2024; Jakovac et al., 2016). Environmental filters such as degraded soils (Poorter et al., 2019), frequent fires (Vedovato et al., 2025), severe droughts (Elias et al., 2020) and distance to old growth undisturbed primary forest (Magnago et al., 2017) may help explain this pattern, particularly in the early stages when only a few species are able to colonize, establish, and dominate in terms of carbon storage and abundance (Boukili & Chazdon, 2017; Lohbeck et al., 2014). However, hyperdominance within the community assembly process is also likely to be an outcome of their high dispersal capacity, with small seeded species dispersed by bats and birds being able to arrive first (González-Castro et al., 2019; Hawes et al., 2020; Wendt et al., 2022).

### 4.2 Palms as hyperdominants in secondary forests

The presence of palms among hyperdominant species, such as *A. speciosa and A. maripa*, provides strong evidence that this group has a significant influence on the Amazonian woody flora. In line with these results, other studies have shown that palms are among the most common species in Amazonian primary forests, with many listed as hyperdominant species, such as *A. maripa* (ter Steege et al., 2013). Their spatial distribution has been linked to various factors, including human presence, high efficiency of germination and dispersal and potential climate “resilience” (Muscarella et al., 2020; Viana et al., 2021; Santos et al., 2025). Their prevalence here matches their known importance in succession (Boukili & Chazdon, 2017), even when their presence has not been intentionally enriched in regrowing forests. The impact of palms on other tree species remains poorly understood, but some studies suggest they can be invasive (Freitas et al., 2021; Gehring et al., 2020) and may impede succession by forming recalcitrant understorey layers (Royo & Carson, 2006). In our plots, this group was among the dominant species in terms of both carbon stocks and abundance – but their role with trees requires investigation. Although palms can dominate, they also have great potential to provide shade and attract pollinators and seed dispersers (Barfod et al., 2011; Lim et al., 2020) that can support more diverse trajectories of natural regeneration.

### 4.3 Ecological stability of hyperdominants contrasts with their low conservation value

Changes in wood density values, a key trait for plant growth, survival and forest carbon storage, reveal two important insights about the hyperdominance process in the secondary forests. First, the hyperdominant species had lower wood densities than non-hyperdominants, with around half above the average (0.68) and values ranging from 0.22 to 0.99 (Figure 5). Furthermore, the species that contributed the most to abundance and carbon storage across the plots, such as *C. palmata*, *A. exsucca and T. guianensis* have wood densities below the average of hyperdominant species. These results mean that secondary forests will have lower carbon storage capacities than old-growth forests even when they have similar structures – a point that is important in studies that use LiDAR and can only observe forest structure. The lower wood density could also be important for determining the sensitivity of these ecosystems to disturbances such as severe droughts (Elias et al., 2020) particularly if wood density reflects water transport, the probability of embolisms (Markesteijn et al., 2011; Pineda-García et al., 2013) and survival rates (Poorter et al., 2010).

Second, the persistence of lower wood density among hyperdominant species over time shows how the successional process will be slow, with a prolonged presence of long-lived pioneer species throughout extended successional periods. Whilst this may not necessarily indicate impeded or arrested succession (Fernandes Neto et al., 2019; Mesquita et al., 2001), it does reflect the long periods taken for community turnover and growth required to restore forests that approximate old-growth conditions (Elias et al., 2020). In addition to preserving hyperdominance patterns among larger individuals (Draper et al., 2021), small-stemmed trees can reflect the forests of the future and provide valuable insights into successional development (Lennox et al., 2018). Assessments of this size class (Figure SI 7) indicate that the long-term trajectory of functional succession will progress over time, with both hyperdominant and non-hyperdominant species exhibiting increases in average wood density as forests recover.

### 4.4 Implications for research and large-scale restoration

Identifying the hyperdominant set of Amazonian species in secondary forests opens up new avenues for research and restoration planning. First, secondary forests are sensitive to climatic stress (Elias et al., 2020), and evaluating the ecophysiology of a relatively limited set of species could help further our understanding of climate sensitivities, adding to the knowledge derived from primary forests (Tavares et al., 2023). Second, remote sensing of canopy traits has enabled large-scale assessment of old-growth forests (Aguirre-Gutiérrez et al., 2025). As these hyperdominant species cover a significant proportion of forest carbon and abundance, they may provide a way of linking forest conditions to remotely sensed variables, such as canopy reflectance.

Finally, the presence of hyperdominant species could provide a quick way to help gauge the ecological integrity of assisted and naturally regenerating secondary forests, which will be important as national initiatives (Brasil, 2024) and Amazonian states such as Pará invest in large-scale restoration (SEMAS, 2020, 2023) and national initiatives. The relatively short list of hyperdominant species such as *C. palmata*, *A. exsucca*, *T. guianensis*, and *I. alba* (see Figure 4) could also help guide assisted restoration and enrichment strategies, providing a clear focus on species that are known to be successful and ecologically important across a broad suite of regeneration contexts. These species are not just better adapted to the spectrum of environmental conditions found in secondary forests; they also act as key species in recovering carbon stocks and forest structure. In doing so, they become the foundation species in forests that harbour a much higher diversity of species, and may play a role in helping these later successional and old growth species, such as *E. coriacea*, *Ormosia paraensis*, *Sacoglottis guianensis* and *Bertholletia excelsa,* to overcome the environmental filtering that exists at the start of the successional process (Lohbeck et al. 2013). The high diversity of non-dominants is likely to be crucial for restoration to be functionally effective and resilient over time, supporting multiple ecosystem functions beyond carbon (Di Sacco et al., 2021; Ferreira et al., 2018) including their long-term resilience to climate change (Arroyo-Rodríguez et al., 2017).

## Data availability statement

All database is available on the ForestPlot.net platform and can be accessed upon request.

## Acknowledgements

We are grateful to the following for financial support: Conselho Nacional de Desenvolvimento Científico e Tecnológico – CNPq (PELD-RAS, Process No. 445994/2024-0, CAPOEIRA, No. CNPq/443849/2024-2), Instituto Nacional de Ciência e Tecnologia – Biodiversidade e Uso da Terra na Amazônia (CNPq 574008/2008-0), Empresa Brasileira de Pesquisa Agropecuária – Embrapa (SEG:02.08.06.005.00), the UK government Darwin Initiative (17-023), The Nature Conservancy, and Natural Environment Research Council (NERC) (NE/F01614X/1, NE/G000816/1, NE/K016431/1, NE/S01084X/1, NE/X015262/1, and NE/X019039/1), the UK government’s Global Centre of Biodiversity for Climate, and the BNP Paribas Foundation’s Biodiversity and Climate Initiative. We are deeply grateful to our numerous field and laboratory assistants. We also thank the farmers and workers unions of Santarém, Belterra, and Paragominas and all collaborating private landowners for their support. We thank the Large-Scale Biosphere-Atmosphere Program (LBA) for logistical and infrastructure support during field measurements in the Santarém region. FE supported by the Serrapilheira Institute fellowship/FAPESPA (grant number – R-2401–46863, TO No. 158/2024). ICGV acknowledge for INCT Nexus grants (CNPq 406516/2022-7and Fapespa E-2024/2215630). This paper is number #129 in the Rede Amazônia Sustentável publication series (https://ras-network.org).

## Author contributions

FE: Conceptualization, Methodology, Data collection, Formal analysis, Writing – review & editing. EB, DVS, RON, ICGV, LM, JF: Writing – review & editing, Data collection. JB and JF: Conceptualization, Writing – review & editing, Supervision and Project administration.

## References

1. Aguirre-Gutiérrez, J., Rifai, S. W., Deng, X., ter Steege, H., Thomson, E., Corral-Rivas, J. J., Guimaraes, A. F., Muller, S., Klipel, J., Fauset, S., Resende, A. F., Wallin, G., Joly, C. A., Abernethy, K., Adu-Bredu, S., Alexandre Silva, C., de Oliveira, E. A., Almeida, D. R. A., Alvarez-Davila, E., … Malhi, Y. (2025). Canopy functional trait variation across Earth’s tropical forests. Nature, 1–8. 10.1038/s41586-025-08663-2

2. Arroyo-Rodríguez, V., Melo, F. P. L., Martínez-Ramos, M., Bongers, F., Chazdon, R. L., Meave, J. A., Norden, N., Santos, B. A., Leal, I. R., & Tabarelli, M. (2017). Multiple successional pathways in human-modified tropical landscapes: New insights from forest succession, forest fragmentation and landscape ecology research: Multiple successional pathways. Biological Reviews, 92(1), Article 1. 10.1111/brv.12231

3. Barfod, A. S., Hagen, M., & Borchsenius, F. (2011). Twenty-five years of progress in understanding pollination mechanisms in palms (Arecaceae). Annals of Botany, 108(8), 1503–1516. 10.1093/aob/mcr192

4. Barlow, J., Sist, P., Almeida, R., Arantes, C. C., Berenguer, E., Caron, P., Cuesta, F., Doria, C., Ferreira, J., Flecker, A., Heilpern, S., Kalamandeen, M., Lees, A. C., & Nasci, N. (2022). Restoration options for the Amazon. In The Amazon We Want.

5. Bennett, A. C., Rodrigues de Sousa, T., Monteagudo-Mendoza, A., Esquivel-Muelbert, A., Morandi, P. S., Coelho de Souza, F., Castro, W., Duque, L. F., Flores Llampazo, G., Manoel dos Santos, R., Ramos, E., Vilanova Torre, E., Alvarez-Davila, E., Baker, T. R., Costa, F. R. C., Lewis, S. L., Marimon, B. S., Schietti, J., Burban, B., … Phillips, O. L. (2023). Sensitivity of South American tropical forests to an extreme climate anomaly. Nature Climate Change, 13(9), Article 9. 10.1038/s41558-023-01776-4

6. BFG. (2018). Brazilian Flora 2020: Innovation and collaboration to meet Target 1 of the Global Strategy for Plant Conservation (GSPC). 69, 1513.

7. Boukili, V. K., & Chazdon, R. L. (2017). Environmental filtering, local site factors and landscape context drive changes in functional trait composition during tropical forest succession. Perspectives in Plant Ecology, Evolution and Systematics, 24, 37–47. 10.1016/j.ppees.2016.11.003

8. Brasil. (2024). Plano Nacional de Recuperação da Vegetação Nativa – Planaveg 2025-2028. https://www.gov.br/mma/pt-br/composicao/sbio/dflo/plano-nacional-de-recuperacao-da-vegetacao-nativa-planaveg/planaveg_2025-2028_2dez2024.pdf

9. Carle, H., Bauman, D., Evans, M. N., Coughlin, I., Binks, O., Ford, A., Bradford, M., Nicotra, A., Murphy, H., & Meir, P. (2025). Aboveground biomass in Australian tropical forests now a net carbon source. Nature, 646(8085), 611–618. 10.1038/s41586-025-09497-8

10. Chave, J., Réjou-Méchain, M., Búrquez, A., Chidumayo, E., Colgan, M. S., Delitti, W. B. C., Duque, A., Eid, T., Fearnside, P. M., Goodman, R. C., Henry, M., Martínez-Yrízar, A., Mugasha, W. A., Muller-Landau, H. C., Mencuccini, M., Nelson, B. W., Ngomanda, A., Nogueira, E. M., Ortiz-Malavassi, E., … Vieilledent, G. (2014). Improved allometric models to estimate the aboveground biomass of tropical trees. Global Change Biology, 20(10), Article 10. 10.1111/gcb.12629

11. Chazdon, R. L. (2014). Second Growth: The Promise of Tropical Forest Regeneration in an Age of Deforestation (Vol. 1). University of Chicago Press. 10.7208/9780226118109

12. Chazdon, R. L., Finegan, B., Capers, R. S., Salgado-Negret, B., Casanoves, F., Boukili, V., & Norden, N. (2010). Composition and Dynamics of Functional Groups of Trees During Tropical Forest Succession in Northeastern Costa Rica: Functional Groups of Trees. Biotropica, 42(1), Article 1. 10.1111/j.1744-7429.2009.00566.x

13. da Silva, C. M., Elias, F., do Nascimento, R. O., & Ferreira, J. (2023). The potential for forest landscape restoration in the Amazon: State of the art of restoration strategies. Restoration Ecology, 31(4), Article 4. 10.1111/rec.13955

14. Di Sacco, A., Hardwick, K. A., Blakesley, D., Brancalion, P. H. S., Breman, E., Cecilio Rebola, L., Chomba, S., Dixon, K., Elliott, S., Ruyonga, G., Shaw, K., Smith, P., Smith, R. J., & Antonelli, A. (2021). Ten golden rules for reforestation to optimize carbon sequestration, biodiversity recovery and livelihood benefits. Global Change Biology, 27(7), Article 7. 10.1111/gcb.15498

15. Draper, F. C., Costa, F. R. C., Arellano, G., Phillips, O. L., Duque, A., Macía, M. J., ter Steege, H., Asner, G. P., Berenguer, E., Schietti, J., Socolar, J. B., de Souza, F. C., Dexter, K. G., Jørgensen, P. M., Tello, J. S., Magnusson, W. E., Baker, T. R., Castilho, C. V., Monteagudo-Mendoza, A., … Baraloto, C. (2021). Amazon tree dominance across forest strata. Nature Ecology & Evolution, 5(6), Article 6. 10.1038/s41559-021-01418-y

16. Elias, F., Ferreira, J., Lennox, G. D., Berenguer, E., Ferreira, S., Schwartz, G., Melo, L. de O., Reis Júnior, D. N., Nascimento, R. O., Ferreira, F. N., Espirito-Santo, F., Smith, C. C., & Barlow, J. (2020). Assessing the growth and climate sensitivity of secondary forests in highly deforested Amazonian landscapes. Ecology, 101(3), Article 3. 10.1002/ecy.2954

17. Elias, F., Ferreira, J., Resende, A. F., Berenguer, E., França, F., Smith, C. C., Schwartz, G., Nascimento, R. O., Guedes, M., Chesini Rossi, L., Maria Moraes de Seixas, M., Melo da Silva, C., & Barlow, J. (2022). Comparing contemporary and lifetime rates of carbon accumulation from secondary forests in the eastern Amazon. Forest Ecology and Management, 508, 120053. 10.1016/j.foreco.2022.120053

18. Esquivel-Muelbert, A., Baker, T. R., Dexter, K. G., Lewis, S. L., Brienen, R. J. W., Feldpausch, T. R., Lloyd, J., Monteagudo-Mendoza, A., Arroyo, L., Álvarez-Dávila, E., Higuchi, N., Marimon, B. S., Marimon-Junior, B. H., Silveira, M., Vilanova, E., Gloor, E., Malhi, Y., Chave, J., Barlow, J., … Phillips, O. L. (2019). Compositional response of Amazon forests to climate change. Global Change Biology, 25(1), Article 1. 10.1111/gcb.14413

19. Fauset, S., Johnson, M. O., Gloor, M., Baker, T. R., Monteagudo M., A., Brienen, R. J. W., Feldpausch, T. R., Lopez-Gonzalez, G., Malhi, Y., ter Steege, H., Pitman, N. C. A., Baraloto, C., Engel, J., Pétronelli, P., Andrade, A., Camargo, J. L. C., Laurance, S. G. W., Laurance, W. F., Chave, J., … Phillips, O. L. (2015). Hyperdominance in Amazonian forest carbon cycling. Nature Communications, 6(1), Article 1. 10.1038/ncomms7857

20. Fernandes Neto, J. G., Costa, F. R. C., Williamson, G. B., & Mesquita, R. C. G. (2019). Alternative functional trajectories along succession after different land uses in central Amazonia. Journal of Applied Ecology, 56(11), Article 11. 10.1111/1365-2664.13484

21. Ferreira, J., Lennox, G. D., Gardner, T. A., Thomson, J. R., Berenguer, E., Lees, A. C., Mac Nally, R., Aragão, L. E. O. C., Ferraz, S. F. B., Louzada, J., Moura, N. G., Oliveira, V. H. F., Pardini, R., Solar, R. R. C., Vieira, I. C. G., & Barlow, J. (2018). Carbon-focused conservation may fail to protect the most biodiverse tropical forests. Nature Climate Change, 8(8), Article 8. 10.1038/s41558-018-0225-7

22. Freitas, M. A. B., Magalhães, J. L. L., Carmona, C. P., Arroyo-Rodríguez, V., Vieira, I. C. G., & Tabarelli, M. (2021). Intensification of açaí palm management largely impoverishes tree assemblages in the Amazon estuarine forest. Biological Conservation, 261, 109251. 10.1016/j.biocon.2021.109251

23. Gardner, T. A., Ferreira, J., Barlow, J., Lees, A. C., Parry, L., Vieira, I. C. G., Berenguer, E., Abramovay, R., Aleixo, A., Andretti, C., Aragão, L. E. O. C., Araújo, I., de Ávila, W. S., Bardgett, R. D., Batistella, M., Begotti, R. A., Beldini, T., de Blas, D. E., Braga, R. F., … Zuanon, J. (2013). A social and ecological assessment of tropical land uses at multiple scales: The Sustainable Amazon Network. Philosophical Transactions of the Royal Society B: Biological Sciences, 368(1619), Article 1619. 10.1098/rstb.2012.0166

24. Garrett, R. D., Cammelli, F., Ferreira, J., Levy, S. A., Valentim, J., & Vieira, I. (2021). Forests and Sustainable Development in the Brazilian Amazon: History, Trends, and Future Prospects. Annual Review of Environment and Resources, 46(1), Article 1. 10.1146/annurev-environ-012220-010228

25. Gehring, C., Zelarayán, M. C., Luz, R. L., Almeida, R. B., Boddey, R. M., & Leite, M. F. A. (2020). Babassu palm (Attalea speciosa Mart.) super-dominance shapes its surroundings via multiple biotic, soil chemical, and physical interactions and accumulates soil carbon: A case study in eastern Amazonia. Plant and Soil, 454(1), 447–460. 10.1007/s11104-020-04580-7

26. Giles, A. L., Schietti, J., Rosenfield, M. F., Mesquita, R. C., Vieira, D. L. M., Vieira, I. C. G., Poorter, L., Brancalion, P. H. S., Peña-Claros, M., Siqueira, J., Oliveira Junior, L., do Espírito-Santo, M. M., Sarmento, P. S. de M., Ferreira, J. N., Berenguer, E., Barlow, J., Elias, F., Cassol, H. L. G., Silva, R. C., … Jakovac, C. C. (2024). Simple ecological indicators benchmark regeneration success of Amazonian forests. Communications Earth & Environment, 5(1), 1–12. 10.1038/s43247-024-01949-9

27. González-Castro, A., Yang, S., & Carlo, T. A. (2019). How does avian seed dispersal shape the structure of early successional tropical forests? Functional Ecology, 33(2), 229–238. 10.1111/1365-2435.13250

28. Goodman, R. C., Phillips, O. L., del Castillo Torres, D., Freitas, L., Cortese, S. T., Monteagudo, A., & Baker, T. R. (2013). Amazon palm biomass and allometry. Forest Ecology and Management, 310, 994–1004. 10.1016/j.foreco.2013.09.045

29. Hawes, J. E., Vieira, I. C. G., Magnago, L. F. S., Berenguer, E., Ferreira, J., Aragão, L. E. O. C., Cardoso, A., Lees, A. C., Lennox, G. D., Tobias, J. A., Waldron, A., & Barlow, J. (2020). A large-scale assessment of plant dispersal mode and seed traits across human-modified Amazonian forests. Journal of Ecology, 108(4), Article 4. 10.1111/1365-2745.13358

30. Jakovac, C. C., Bongers, F., Kuyper, T. W., Mesquita, R. C. G., & Peña-Claros, M. (2016). Land use as a filter for species composition in Amazonian secondary forests. Journal of Vegetation Science, 27(6), 1104–1116. 10.1111/jvs.12457

31. Jakovac, C. C., Junqueira, A. B., Crouzeilles, R., Peña-Claros, M., Mesquita, R. C. G., & Bongers, F. (2021). The role of land-use history in driving successional pathways and its implications for the restoration of tropical forests. Biological Reviews, 96(4), Article 4. 10.1111/brv.12694

32. Jakovac, C. C., Meave, J. A., Bongers, F., Letcher, S. G., Dupuy, J. M., Piotto, D., Rozendaal, D. M. A., Peña-Claros, M., Craven, D., Santos, B. A., Siminski, A., Fantini, A. C., Rodrigues, A. C., Hernández-Jaramillo, A., Idárraga, A., Junqueira, A. B., Zambrano, A. M. A., de Jong, B. H. J., Pinho, B. X., … Poorter, L. (2022). Strong floristic distinctiveness across Neotropical successional forests. Science Advances, 8(26), Article 26. 10.1126/sciadv.abn1767

33. Lapola, D. M., Pinho, P., Barlow, J., Aragão, L. E. O. C., Berenguer, E., Carmenta, R., Liddy, H. M., Seixas, H., Silva, C. V. J., Silva-Junior, C. H. L., Alencar, A. A. C., Anderson, L. O., Armenteras, D., Brovkin, V., Calders, K., Chambers, J., Chini, L., Costa, M. H., Faria, B. L., … Walker, W. S. (2023). The drivers and impacts of Amazon forest degradation. Science, 379(6630), Article 6630. 10.1126/science.abp8622

34. Lennox, G. D., Gardner, T. A., Thomson, J. R., Ferreira, J., Berenguer, E., Lees, A. C., Mac Nally, R., Aragão, L. E. O. C., Ferraz, S. F. B., Louzada, J., Moura, N. G., Oliveira, V. H. F., Pardini, R., Solar, R. R. C., Vaz-de Mello, F. Z., Vieira, I. C. G., & Barlow, J. (2018). Second rate or a second chance? Assessing biomass and biodiversity recovery in regenerating Amazonian forests. Global Change Biology, 24(12), Article 12. 10.1111/gcb.14443

35. Lim, J. Y., Svenning, J.-C., Göldel, B., Faurby, S., & Kissling, W. D. (2020). Frugivore-fruit size relationships between palms and mammals reveal past and future defaunation impacts. Nature Communications, 11(1), 4904. 10.1038/s41467-020-18530-5

36. Lohbeck, M., Poorter, L., Martínez-Ramos, M., Rodriguez-Velázquez, J., van Breugel, M., & Bongers, F. (2014). Changing drivers of species dominance during tropical forest succession. Functional Ecology, 28(4), 1052–1058. 10.1111/1365-2435.12240

37. Lopez-Gonzalez, G., Lewis, S. L., Burkitt, M., & Phillips, O. L. (2011). ForestPlots.net: A web application and research tool to manage and analyse tropical forest plot data: ForestPlots.net. Journal of Vegetation Science, 22(4), Article 4. 10.1111/j.1654-1103.2011.01312.x

38. Magnago, L. F. S., Magrach, A., Barlow, J., Schaefer, C. E. G. R., Laurance, W. F., Martins, S. V., & Edwards, D. P. (2017). Do fragment size and edge effects predict carbon stocks in trees and lianas in tropical forests? Functional Ecology, 31(2), 542–552. 10.1111/1365-2435.12752

39. Markesteijn, L., Poorter, L., Paz, H., Sack, L., & Bongers, F. (2011). Ecological differentiation in xylem cavitation resistance is associated with stem and leaf structural traits. Plant, Cell & Environment, 34(1), 137–148. 10.1111/j.1365-3040.2010.02231.x

40. Mesquita, R. C. G., Ickes, K., Ganade, G., & Williamson, G. B. (2001). Alternative successional pathways in the Amazon Basin. Journal of Ecology, 89(4), Article 4. 10.1046/j.1365-2745.2001.00583.x

41. Muscarella, R., Emilio, T., Phillips, O. L., Lewis, S. L., Slik, F., Baker, W. J., Couvreur, T. L. P., Eiserhardt, W. L., Svenning, J.-C., Affum-Baffoe, K., Aiba, S.-I., de Almeida, E. C., de Almeida, S. S., de Oliveira, E. A., Álvarez-Dávila, E., Alves, L. F., Alvez-Valles, C. M., Carvalho, F. A., Guarin, F. A., … Balslev, H. (2020). The global abundance of tree palms. Global Ecology and Biogeography, 29(9), Article 9. 10.1111/geb.13123

42. Nelson, B. W., Mesquita, R., Pereira, J. L. G., Garcia Aquino de Souza, S., Teixeira Batista, G., & Bovino Couto, L. (1999). Allometric regressions for improved estimate of secondary forest biomass in the central Amazon. Forest Ecology and Management, 117(1), 149–167. 10.1016/S0378-1127(98)00475-7

43. Nunes, S., Oliveira, L., Siqueira, J., Morton, D. C., & Souza, C. M. (2020). Unmasking secondary vegetation dynamics in the Brazilian Amazon. Environmental Research Letters, 15(3), Article 3. 10.1088/1748-9326/ab76db

44. Pereira, M. B., Elias, F., Teixeira, N. D. A., Feldpausch, T. R., Marimon-Junior, B. H., & Marimon, B. S. (2025). Post-fire changes in tree diversity, composition and carbon in seasonal forests in the Southern Amazonia. Forest Ecology and Management, 578, 122447. 10.1016/j.foreco.2024.122447

45. Pineda-García, F., Paz, H., & Meinzer, F. C. (2013). Drought resistance in early and late secondary successional species from a tropical dry forest: The interplay between xylem resistance to embolism, sapwood water storage and leaf shedding. Plant, Cell & Environment, 36(2), 405–418. 10.1111/j.1365-3040.2012.02582.x

46. Pinho, B. X., Melo, F. P. L., ter Braak, C. J. F., Bauman, D., Maréchaux, I., Tabarelli, M., Benchimol, M., Arroyo-Rodriguez, V., Santos, B. A., Hawes, J. E., Berenguer, E., Ferreira, J., Silveira, J. M., Peres, C. A., Rocha-Santos, L., Souza, F. C., Gonçalves-Souza, T., Mariano-Neto, E., Faria, D., & Barlow, J. (2024). Winner–loser plant trait replacements in human-modified tropical forests. Nature Ecology & Evolution, 1–15. 10.1038/s41559-024-02592-5

47. Poorter, L., Amissah, L., Bongers, F., Hordijk, I., Kok, J., Laurance, S. G. W., Lohbeck, M., Martínez-Ramos, M., Matsuo, T., Meave, J. A., Muñoz, R., Peña-Claros, M., & van der Sande, M. T. (2023). Successional theories. Biological Reviews, 98(6), Article 6. 10.1111/brv.12995

48. Poorter, L., Bongers, F., Aide, T. M., Almeyda Zambrano, A. M., Balvanera, P., Becknell, J. M., Boukili, V., Brancalion, P. H. S., Broadbent, E. N., Chazdon, R. L., Craven, D., de Almeida-Cortez, J. S., Cabral, G. A. L., de Jong, B. H. J., Denslow, J. S., Dent, D. H., DeWalt, S. J., Dupuy, J. M., Durán, S. M., … Rozendaal, D. M. A. (2016). Biomass resilience of Neotropical secondary forests. Nature, 530(7589), Article 7589. 10.1038/nature16512

49. Poorter, L., McDonald, I., Alarcón, A., Fichtler, E., Licona, J.-C., Peña-Claros, M., Sterck, F., Villegas, Z., & Sass-Klaassen, U. (2010). The importance of wood traits and hydraulic conductance for the performance and life history strategies of 42 rainforest tree species. New Phytologist, 185(2), 481–492. 10.1111/j.1469-8137.2009.03092.x

50. Poorter, L., Rozendaal, D. M. A., Bongers, F., de Almeida-Cortez, J. S., Almeyda Zambrano, A. M., Álvarez, F. S., Andrade, J. L., Villa, L. F. A., Balvanera, P., Becknell, J. M., Bentos, T. V., Bhaskar, R., Boukili, V., Brancalion, P. H. S., Broadbent, E. N., César, R. G., Chave, J., Chazdon, R. L., Colletta, G. D., … Westoby, M. (2019). Wet and dry tropical forests show opposite successional pathways in wood density but converge over time. Nature Ecology & Evolution, 3(6), Article 6. 10.1038/s41559-019-0882-6

51. R Core Team. (2025). R: A Language and Environment for Statistical Computing. https://www.r-project.org/

52. Royo, A. A., & Carson, W. P. (2006). On the formation of dense understory layers in forests worldwide: Consequences and implications for forest dynamics, biodiversity, and succession. Canadian Journal of Forest Research, 36(6), 1345–1362. 10.1139/x06-025

53. Santos, D. P., Silva, T. S. F., & de Assis Figueiredo, F. A. M. M. (2025). Climate Change May Increase Environmental Suitability of the Babassu Complex (Attalea spp., Arecaceae). Journal of Biogeography, 52(11), e70027. 10.1111/jbi.70027

54. SEMAS. (2020, August 3). Plano Estadual Amazônia Agora, decreto n° 941. https://www.semas.pa.gov.br/wp-content/uploads/2021/02/GUIAINFO.pdf

55. SEMAS. (2023). Plano de Recuperação da Vegetação Nativa do Estado do Pará (PRVN-PA). https://semas.pa.gov.br/prvn/

56. SEMAS. (2024). Plano Estadual de Bioeconomia do Pará. Secretaria de Estado de Meio Ambiente e Sustentabilidade do Estado do Pará. https://www.semas.pa.gov.br/planbio

57. Smith, C. C., Barlow, J., Healey, J. R., Miranda, L. de S., Young, P. J., & Schwartz, N. B. (2023). Amazonian secondary forests are greatly reducing fragmentation and edge exposure in old-growth forests. Environmental Research Letters, 18(12), Article 12. 10.1088/1748-9326/ad039e

58. Smith, C. C., Healey, J. R., Berenguer, E., Young, P. J., Taylor, B., Elias, F., Espírito-Santo, F., & Barlow, J. (2021). Old-growth forest loss and secondary forest recovery across Amazonian countries. Environmental Research Letters, 16(8), Article 8. 10.1088/1748-9326/ac1701

59. Solar, R. R. de C., Barlow, J., Ferreira, J., Berenguer, E., Lees, A. C., Thomson, J. R., Louzada, J., Maués, M., Moura, N. G., Oliveira, V. H. F., Chaul, J. C. M., Schoereder, J. H., Vieira, I. C. G., Mac Nally, R., & Gardner, T. A. (2015). How pervasive is biotic homogenization in human-modified tropical forest landscapes? Ecology Letters, 18(10), Article 10. 10.1111/ele.12494

60. Tavares, J. V., Oliveira, R. S., Mencuccini, M., Signori-Müller, C., Pereira, L., Diniz, F. C., Gilpin, M., Marca Zevallos, M. J., Salas Yupayccana, C. A., Acosta, M., Pérez Mullisaca, F. M., Barros, F. de V., Bittencourt, P., Jancoski, H., Scalon, M. C., Marimon, B. S., Oliveras Menor, I., Marimon, B. H., Fancourt, M., … Galbraith, D. R. (2023). Basin-wide variation in tree hydraulic safety margins predicts the carbon balance of Amazon forests. Nature, 617(7959), Article 7959. 10.1038/s41586-023-05971-3

61. ter Steege, H., Pitman, N. C. A., Sabatier, D., Baraloto, C., Salomão, R. P., Guevara, J. E., Phillips, O. L., Castilho, C. V., Magnusson, W. E., Molino, J.-F., Monteagudo, A., Núñez Vargas, P., Montero, J. C., Feldpausch, T. R., Coronado, E. N. H., Killeen, T. J., Mostacedo, B., Vasquez, R., Assis, R. L., … Silman, M. R. (2013). Hyperdominance in the Amazonian Tree Flora. Science, 342(6156), 1243092. 10.1126/science.1243092

62. Vedovato, L. B., Aragão, L. E. O. C., Almeida, D. R. A., Bartholomew, D. C., Assis, M., Dalagnol, R., Gorgens, E. B., Silva-Junior, C. H. L., Ometto, J. P., Pontes-Lopes, A., Silva, C. A., Valbuena, R., & Feldpausch, T. R. (2025). Impacts of fire on canopy structure and its resilience depend on successional stage in Amazonian secondary forests. Remote Sensing in Ecology and Conservation, 11(4), 394–410. 10.1002/rse2.431

63. Viana, J. L., Turner, B. L., & Dalling, J. W. (2021). Compositional variation in understorey fern and palm communities along a soil fertility and rainfall gradient in a lower montane tropical forest. Journal of Vegetation Science, 32(1), Article 1. 10.1111/jvs.12947

64. Vieira, I. C. G., Giles, A., do Espírito Santo, M. M., Mesquita, R. C. G., Vieira, D. L. M., Massoca, P., Rosenfield, M. F., Albernaz, A. L. M., de Almeida, D. R. A., Vieira, G., Schietti, J., Ferreira, M., Brancalion, P. H. S., & Jakovac, C. C. (2025). Governance and policy constraints of natural forest regeneration in the Brazilian Amazon. Restoration Ecology, 33(1), e14272. 10.1111/rec.14272

65. Wendt, A. L., Chazdon, R. L., & Vargas Ramirez, O. (2022). Successional trajectories of seed dispersal mode and seed size of canopy tree species in wet tropical forests. Frontiers in Forests and Global Change, 5. 10.3389/ffgc.2022.946541

66. Wieland, L. M., Mesquita, R. C. G., Bobrowiec, P. E. D., Bentos, T. V., & Williamson, G. B. (2011). Seed Rain and Advance Regeneration in Secondary Succession in the Brazilian Amazon. Tropical Conservation Science, 4(3), 300–316. 10.1177/194008291100400308

67. Zanne, A. E., Lopez-Gonzalez, G., Coomes, D. A., Ilic, J., Jansen, S., Lewis, S. L., Miller, R. B., Swenson, N. G., Wiemann, M. C., & Chave, J. (2009). *Data from: Towards a worldwide wood economics spectrum* (Version 5, p. 2047488 bytes) [Dataset]. Dryad. 10.5061/DRYAD.234

