## Supporting information for "Hyperdominance and rarity in Amazonian secondary forests"

Running title: **Hyperdominance in Amazonian secondary forests**

Fernando Elias[1*], Joice Ferreira[2], Erika Berenguer[3,4], Rodrigo Oliveira do Nascimento[2], Leonardo Miranda[3], Divino Vicente Silvério[1], Ima Célia Guimarães Vieira[5], Luiz E. O. Aragão[6], Gustavo Schwartz[2], Jos Barlow[3]

1 Universidade Federal Rural da Amazônia, Capitão Poço, Pará, Brazil.

2 Embrapa Amazônia Oriental, Belém, Pará, Brazil.

3 Lancaster Environment Centre, Lancaster University, Lancaster, UK

4 Environmental Change Institute, School of Geography and the Environment, University of Oxford, Oxford, UK.

5 Museu Paraense Emílio Goeldi, Belém, Pará, Brazil

6 Instituto Nacional de Pesquisas Espaciais, São José dos Campos, Brazil

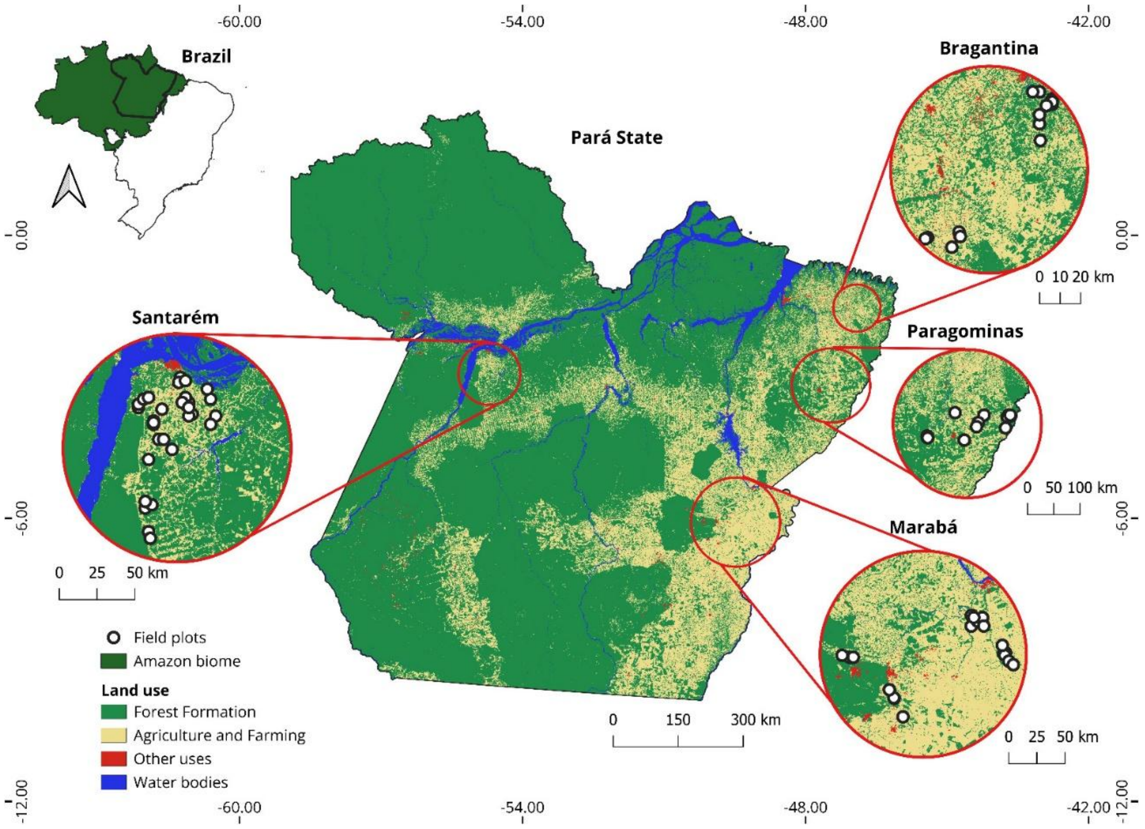


Figure SI 1. Map showing the distribution of field plots and the land use and coverage of the evaluated regions in eastern Amazon. Fonte: Collection 9.0 (MapBiomas, 2024).

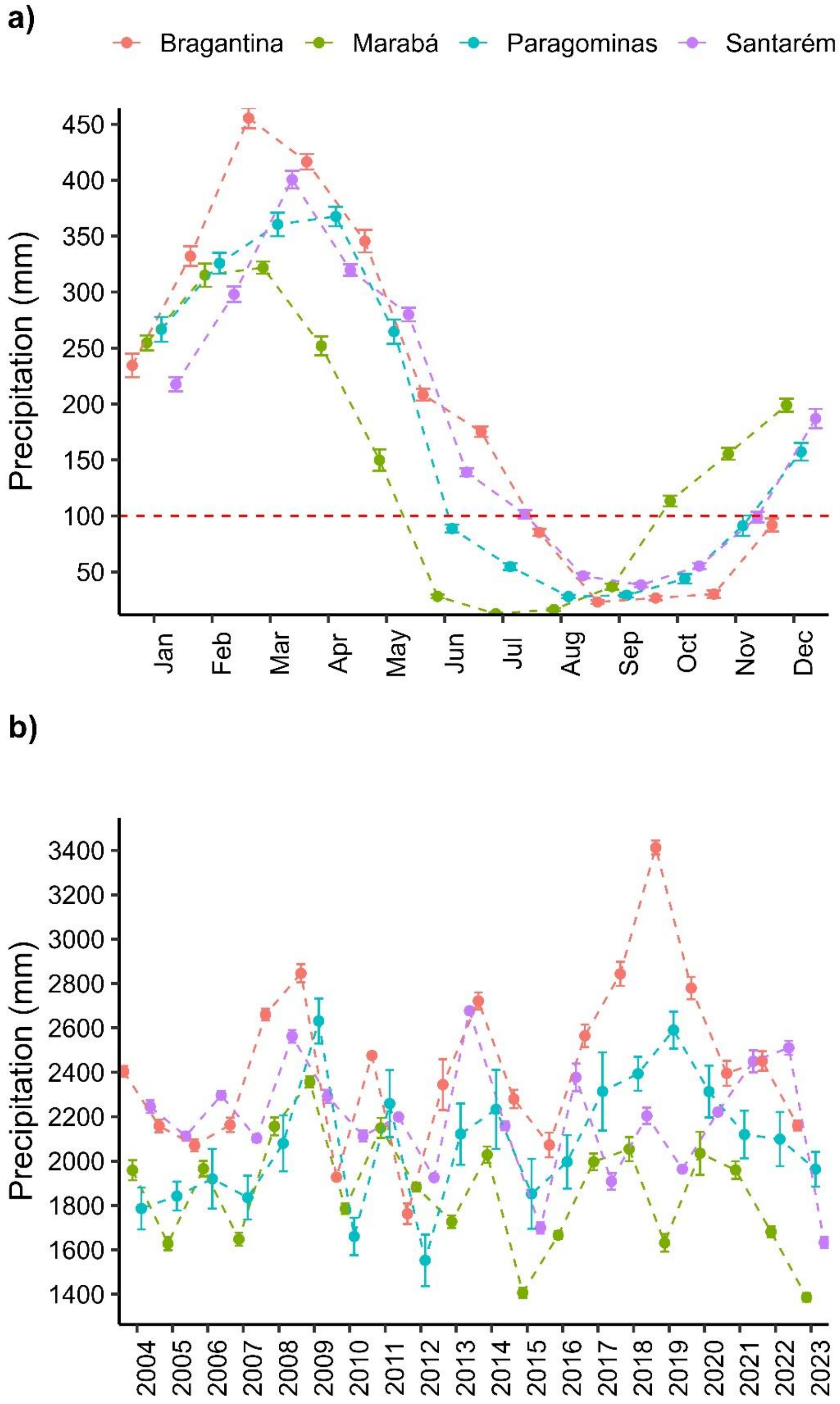


Figure SI 2. Means and confidence intervals of precipitation in secondary forest plots across different sampled regions from 2004 to 2023. Panel A shows the average monthly variation, while Panel B presents the annual variation. Data were extracted from CHIRPS database (Funk et al., 2015).

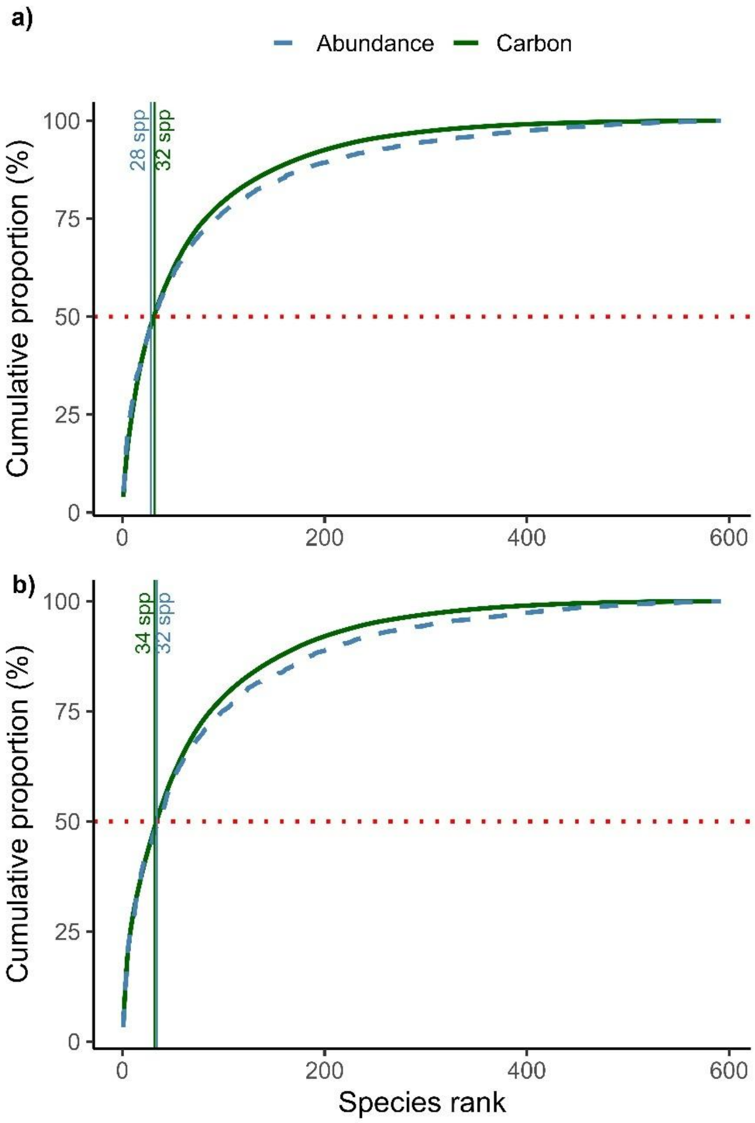


Figure SI 3. Rank of species dominating carbon storage and abundance of small stems. Panel A shows unweighted hyperdominance based on cumulative values across all plots; Panel B shows plot-level weighted hyperdominance. The dashed line marks the species contributing to 50% of cumulative dominance. Vertical lines indicate hyperdominant species identified by each approach and parameter.

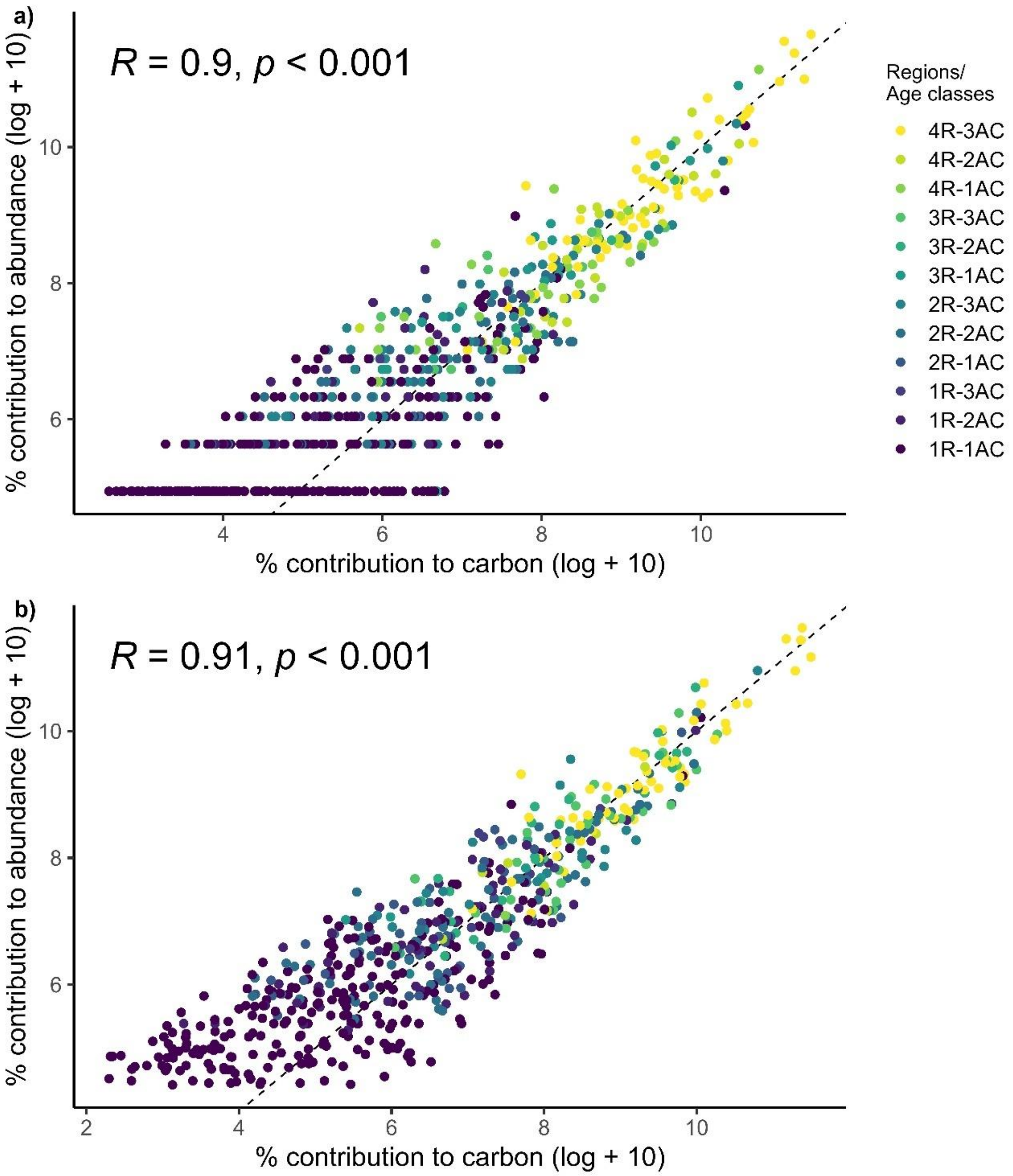


Figure SI 4. Relationship between the relative contributions to abundance and carbon for small stems based on (a) unweighted and (b) plot-weighted approaches. Both axes show log-transformed values (log + 10). The black line represents the trend in the relationship.

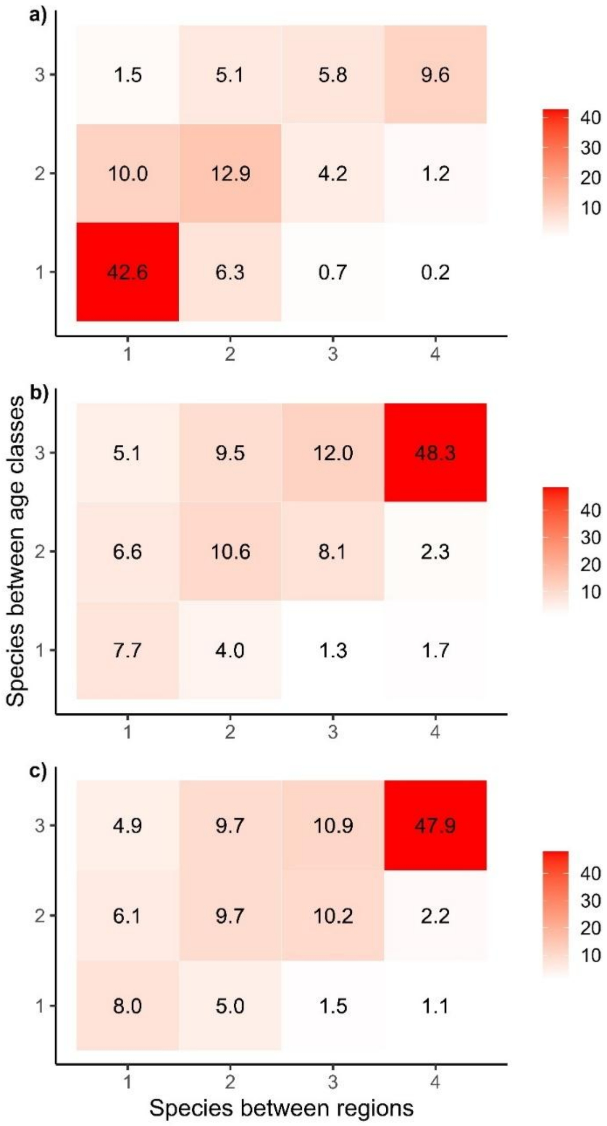


Figure SI 5. Hyperdominance among smaller stems of woody species in Amazonian secondary forests. Panel (a) shows the proportion of species shared among the four evaluated regions and three forest age classes. Panels (b) and (c) show the relative contributions of carbon and abundance, respectively, for the species present in each interaction shown in panel (a).

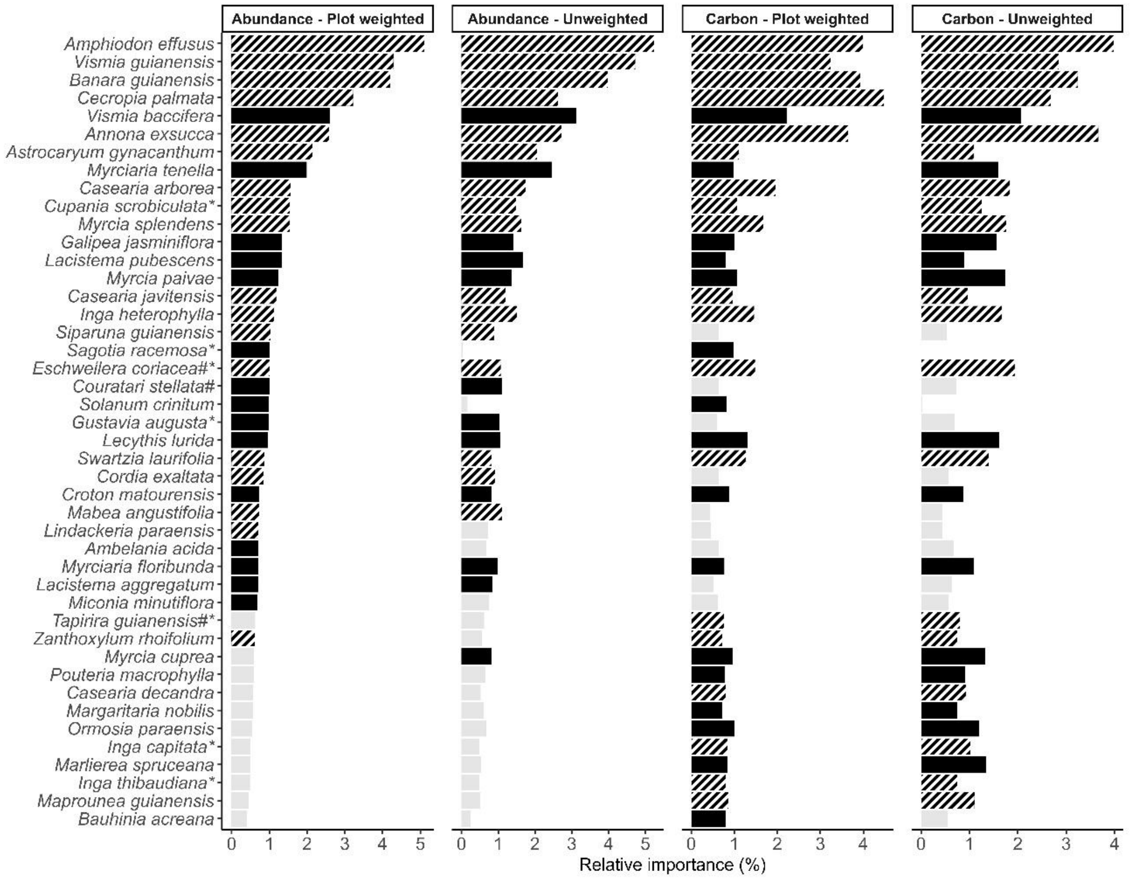


Figure SI 6. Relative contribution of hyperdominant species to carbon storage and abundance based on unweighted and plot-weighted approaches for small stems. Black bars indicate that are hyperdominant species for carbon storage and abundance under both approaches. Black striped bars indicate species present in all four regions and age classes. Light grey bars correspond to non-hyperdominant species. Species are ordered by plot-weighted abundance. * = species classified as abundance hyperdominant by ter Steege et al. (2013); and # = species classified as carbon hyperdominant by Fauset et al. (2015).

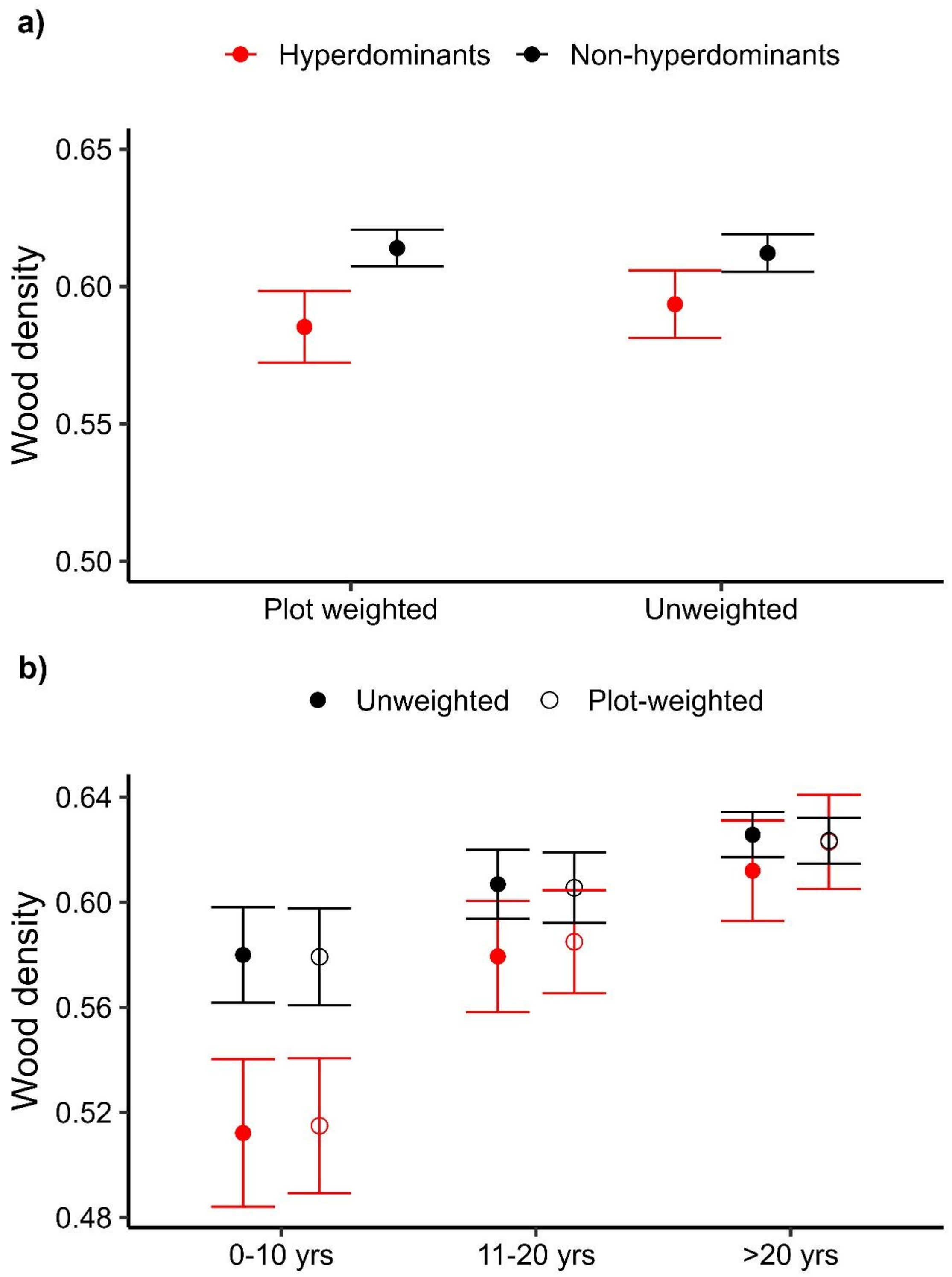


Figure SI 7. Variation in wood density among hiperdominant and non-hyperdominant species among small stems. Panel A shows differences between the two groups across both classification approaches, while panel B shows their variation across forest age classes and both approaches.

**TABLES**

Table SI 1. GAM model estimates of the relationship between height and diameter of trees and palms in secondary forests of eastern Amazonia.

| | **Estimates** | **Std. Error** | **t** | **p-value** | **Habit** |
|---|---|---|---|---|---|
| ~ Diameter | 8.65973 | 0.0169 | 511.8 | 0.000000 | trees |
| ~ Diameter | 10.5763 | 0.1288 | 82.12 | 0.000000 | palms |
| Trees, $R^2$ adj. = 0.741, Scale est. = 7.45, n = 26,033 | | | | | |
| Palms, $R^2$ adj. = 0.664, Scale est. = 8.79, n = 530 | | | | | |

Table SI 2. List of hyperdominant species (both larger and small stems) in our secondary forest plots of the eastern Amazon based on the unweighted and plot weighted approaches. The table shows the cumulative proportions of carbon and abundance for species that together account for 50% of each parameter across all evaluated plots. Species are ordered by cumulative abundance.

| **Family** | **Species** | **Approach** | **Stem size** | **Carbon (%)** | **Abundance (%)** | **No. age classes** | **No. regions** |
|---|---|---|---|---|---|---|---|
| Annonaceae | *Annona exsucca* | Unweighted | Larger | 4.0 | 6.3 | 3 | 4 |
| Urticaceae | *Cecropia palmata* | Unweighted | Larger | 1.3 | 4.6 | 3 | 4 |
| Anacardiaceae | *Tapirira guianensis* | Unweighted | Larger | 5.0 | 4.3 | 3 | 4 |
| Fabaceae | *Inga alba* | Unweighted | Larger | 6.5 | 3.4 | 3 | 4 |
| Bignoniaceae | *Jacaranda copaia* | Unweighted | Larger | 4.0 | 3.1 | 3 | 4 |
| Euphorbiaceae | *Croton matourensis* | Unweighted | Larger | 2.1 | 2.5 | 2 | 3 |
| Euphorbiaceae | *Maprounea guianensis* | Unweighted | Larger | 2.0 | 2.1 | 2 | 3 |
| Arecaceae | *Attalea maripa* | Unweighted | Larger | 4.3 | 2.0 | 3 | 4 |
| Melastomataceae | *Bellucia grossularioides* | Unweighted | Larger | 1.9 | 1.9 | 3 | 3 |
| Fabaceae | *Cassia fastuosa* | Unweighted | Larger | 3.1 | 1.6 | 3 | 4 |
| Arecaceae | *Attalea speciosa* | Unweighted | Larger | 7.4 | 1.5 | 3 | 2 |
| Fabaceae | *Ormosia paraensis* | Unweighted | Larger | | 1.5 | 3 | 4 |
| Salicaceae | *Casearia arborea* | Unweighted | Larger | | 1.5 | 3 | 4 |
| Fabaceae | *Inga edulis* | Unweighted | Larger | 1.3 | 1.5 | 3 | 4 |
| Hypericaceae | *Vismia guianensis* | Unweighted | Larger | | 1.4 | 3 | 4 |
| Annonaceae | *Guatteria punctata* | Unweighted | Larger | | 1.4 | 3 | 4 |
| Humiriaceae | *Sacoglottis guianensis* | Unweighted | Larger | 1.8 | 1.3 | 3 | 3 |
| Boraginaceae | *Cordia exaltata* | Unweighted | Larger | | 1.3 | 3 | 4 |
| Fabaceae | *Inga heterophylla* | Unweighted | Larger | | 1.2 | 3 | 4 |

| | | | | | | | |
|---|---|---|---|---|---|---|---|
| Apocynaceae | *Himatanthus articulatus* | Unweighted | Larger | | 1.2 | 3 | 4 |
| Lecythidaceae | *Eschweilera coriacea* | Unweighted | Larger | | 1.2 | 2 | 3 |
| Fabaceae | *Amphiodon effusus* | Unweighted | Larger | | 1.2 | 3 | 4 |
| Urticaceae | *Cecropia distachya* | Unweighted | Larger | | 1.1 | 3 | 4 |
| Anacardiaceae | *Spondias mombin* | Unweighted | Larger | 1.5 | 1.0 | 3 | 2 |
| Salicaceae | *Banara guianensis* | Unweighted | Larger | | 1.0 | 3 | 4 |
| Araliaceae | *Didymopanax morototoni* | Unweighted | Larger | 2.0 | | 3 | 4 |
| Lecythidaceae | *Bertholletia excelsa* | Unweighted | Larger | 2.5 | | 3 | 2 |
| Fabaceae | *Amphiodon effusus* | Unweighted | Small | 4.0 | 5.2 | 3 | 4 |
| Hypericaceae | *Vismia guianensis* | Unweighted | Small | 2.8 | 4.7 | 3 | 4 |
| Salicaceae | *Banara guianensis* | Unweighted | Small | 3.2 | 4.0 | 3 | 4 |
| Hypericaceae | *Vismia baccifera* | Unweighted | Small | 2.1 | 3.1 | 3 | 2 |
| Annonaceae | *Annona exsucca* | Unweighted | Small | 3.7 | 2.7 | 3 | 4 |
| Urticaceae | *Cecropia palmata* | Unweighted | Small | 2.7 | 2.6 | 3 | 4 |
| Myrtaceae | *Myrciaria tenella* | Unweighted | Small | 1.6 | 2.5 | 2 | 3 |
| Arecaceae | *Astrocaryum gynacanthum* | Unweighted | Small | 1.1 | 2.1 | 3 | 4 |
| Salicaceae | *Casearia arborea* | Unweighted | Small | 1.8 | 1.7 | 3 | 4 |
| Lacistemataceae | *Lacistema pubescens* | Unweighted | Small | 0.9 | 1.7 | 3 | 3 |
| Myrtaceae | *Myrcia splendens* | Unweighted | Small | 1.8 | 1.6 | 3 | 4 |
| Fabaceae | *Inga heterophylla* | Unweighted | Small | 1.7 | 1.5 | 3 | 4 |
| Sapindaceae | *Cupania scrobiculata* | Unweighted | Small | 1.3 | 1.5 | 3 | 4 |
| Rutaceae | *Galipea jasminiflora* | Unweighted | Small | 1.6 | 1.4 | 2 | 2 |
| Myrtaceae | *Myrcia paivae* | Unweighted | Small | 1.7 | 1.4 | 1 | 1 |
| Salicaceae | *Casearia javitensis* | Unweighted | Small | 1.0 | 1.2 | 3 | 4 |
| Euphorbiaceae | *Mabea angustifolia* | Unweighted | Small | | 1.1 | 3 | 4 |
| Lecythidaceae | *Couratari stellata* | Unweighted | Small | | 1.1 | 3 | 2 |

| | | | | | | | |
|---|---|---|---|---|---|---|---|
| Lecythidaceae | *Eschweilera coriacea* | Unweighted | Small | 1.9 | 1.1 | 3 | 4 |
| Lecythidaceae | *Lecythis lurida* | Unweighted | Small | 1.6 | 1.0 | 3 | 3 |
| Lecythidaceae | *Gustavia augusta* | Unweighted | Small | | 1.0 | 2 | 3 |
| Myrtaceae | *Myrciaria floribunda* | Unweighted | Small | 1.1 | 1.0 | 2 | 3 |
| Boraginaceae | *Cordia exaltata* | Unweighted | Small | | 0.9 | 3 | 4 |
| Siparunaceae | *Siparuna guianensis* | Unweighted | Small | | 0.9 | 3 | 4 |
| Lacistematacea e | *Lacistema aggregatum* | Unweighted | Small | | 0.8 | 3 | 3 |
| Fabaceae | *Swartzia laurifolia* | Unweighted | Small | 1.4 | 0.8 | 3 | 4 |
| Euphorbiaceae | *Croton matourensis* | Unweighted | Small | 0.9 | 0.8 | 2 | 3 |
| Myrtaceae | *Myrcia cuprea* | Unweighted | Small | 1.3 | 0.8 | 2 | 2 |
| Fabaceae | *Ormosia paraensis* | Unweighted | Small | 1.2 | | 3 | 3 |
| Sapotaceae | *Pouteria macrophylla* | Unweighted | Small | 0.9 | | 3 | 3 |
| Anacardiaceae | *Tapirira guianensis* | Unweighted | Small | 0.8 | | 3 | 4 |
| Phyllanthaceae | *Margaritaria nobilis* | Unweighted | Small | 0.8 | | 2 | 3 |
| Rutaceae | *Zanthoxylum rhoifolium* | Unweighted | Small | 0.7 | | 3 | 4 |
| Myrtaceae | *Marlierea spruceana* | Unweighted | Small | 1.3 | | 1 | 1 |
| Salicaceae | *Casearia decandra* | Unweighted | Small | 0.9 | | 3 | 4 |
| Euphorbiaceae | *Maprounea guianensis* | Unweighted | Small | 1.1 | | 3 | 4 |
| Fabaceae | *Inga thibaudiana* | Unweighted | Small | 0.8 | | 3 | 4 |
| Fabaceae | *Inga capitata* | Unweighted | Small | 1.0 | | 3 | 4 |
| Urticaceae | *Cecropia palmata* | Plot-weighted | Larger | 6.5 | 7.9 | 3 | 4 |
| Annonaceae | *Annona exsucca* | Plot-weighted | Larger | 5.2 | 6.8 | 3 | 4 |
| Anacardiaceae | *Tapirira guianensis* | Plot-weighted | Larger | 4.4 | 3.4 | 3 | 4 |
| Fabaceae | *Inga alba* | Plot-weighted | Larger | 5.1 | 2.9 | 3 | 4 |

| | | | | | | | |
|---|---|---|---|---|---|---|---|
| Arecaceae | *Attalea maripa* | Plot-weighted | Larger | 4.8 | 2.4 | 3 | 4 |
| Euphorbiaceae | *Croton matourensis* | Plot-weighted | Larger | 2.1 | 2.2 | 2 | 3 |
| Hypericaceae | *Vismia guianensis* | Plot-weighted | Larger | 1.2 | 2.1 | 3 | 4 |
| Bignoniaceae | *Jacaranda copaia* | Plot-weighted | Larger | 2.4 | 2.1 | 3 | 4 |
| Fabaceae | *Cassia fastuosa* | Plot-weighted | Larger | 2.9 | 1.8 | 3 | 4 |
| Euphorbiaceae | *Maprounea guianensis* | Plot-weighted | Larger | 1.5 | 1.7 | 2 | 3 |
| Melastomataceae | *Bellucia grossularioides* | Plot-weighted | Larger | 1.6 | 1.5 | 3 | 3 |
| Arecaceae | *Attalea speciosa* | Plot-weighted | Larger | 3.8 | 1.5 | 3 | 2 |
| Fabaceae | *Inga edulis* | Plot-weighted | Larger | 1.5 | 1.4 | 3 | 4 |
| Burseraceae | *Trattinnickia rhoifolia* | Plot-weighted | Larger | 1.3 | 1.4 | 3 | 4 |
| Fabaceae | *Amphiodon effusus* | Plot-weighted | Larger | | 1.3 | 3 | 4 |
| Fabaceae | *Ormosia paraensis* | Plot-weighted | Larger | | 1.3 | 3 | 4 |
| Salicaceae | *Casearia arborea* | Plot-weighted | Larger | | 1.2 | 3 | 4 |
| Salicaceae | *Banara guianensis* | Plot-weighted | Larger | | 1.2 | 3 | 4 |
| Annonaceae | *Guatteria punctata* | Plot-weighted | Larger | | 1.2 | 3 | 4 |

| | | | | | | | |
|---|---|---|---|---|---|---|---|
| Apocynaceae | *Himatanthus articulatus* | Plot-weighted | Larger | 1.3 | 1.2 | 3 | 4 |
| Humiriaceae | *Sacoglottis guianensis* | Plot-weighted | Larger | 1.4 | 1.1 | 3 | 3 |
| Fabaceae | *Inga heterophylla* | Plot-weighted | Larger | | 1.1 | 3 | 4 |
| Fabaceae | *Senegalia polyphylla* | Plot-weighted | Larger | | 1.0 | 3 | 2 |
| Boraginaceae | *Cordia exaltata* | Plot-weighted | Larger | | 1.0 | 3 | 4 |
| Araliaceae | *Didymopanax morototoni* | Plot-weighted | Larger | 1.5 | | 3 | 4 |
| Lecythidaceae | *Bertholletia excelsa* | Plot-weighted | Larger | 1.9 | | 3 | 2 |
| Fabaceae | *Amphiodon effusus* | Plot-weighted | Small | 4.0 | 5.1 | 3 | 4 |
| Hypericaceae | *Vismia guianensis* | Plot-weighted | Small | 3.2 | 4.3 | 3 | 4 |
| Salicaceae | *Banara guianensis* | Plot-weighted | Small | 3.9 | 4.2 | 3 | 4 |
| Urticaceae | *Cecropia palmata* | Plot-weighted | Small | 4.5 | 3.2 | 3 | 4 |
| Hypericaceae | *Vismia baccifera* | Plot-weighted | Small | 2.2 | 2.6 | 3 | 2 |
| Annonaceae | *Annona exsucca* | Plot-weighted | Small | 3.6 | 2.6 | 3 | 4 |
| Arecaceae | *Astrocaryum gynacanthum* | Plot-weighted | Small | 1.1 | 2.1 | 3 | 4 |
| Myrtaceae | *Myrciaria tenella* | Plot-weighted | Small | 1.0 | 2.0 | 2 | 3 |

| | | | | | | | |
|---|---|---|---|---|---|---|---|
| Salicaceae | *Casearia arborea* | Plot-weighted | Small | 1.9 | 1.6 | 3 | 4 |
| Sapindaceae | *Cupania scrobiculata* | Plot-weighted | Small | 1.1 | 1.5 | 3 | 4 |
| Myrtaceae | *Myrcia splendens* | Plot-weighted | Small | 1.7 | 1.5 | 3 | 4 |
| Rutaceae | *Galipea jasminiflora* | Plot-weighted | Small | 1.0 | 1.3 | 2 | 2 |
| Lacistemataceae | *Lacistema pubescens* | Plot-weighted | Small | 0.8 | 1.3 | 3 | 3 |
| Myrtaceae | *Myrcia paivae* | Plot-weighted | Small | 1.1 | 1.2 | 1 | 1 |
| Salicaceae | *Casearia javitensis* | Plot-weighted | Small | 1.0 | 1.2 | 3 | 4 |
| Fabaceae | *Inga heterophylla* | Plot-weighted | Small | 1.5 | 1.1 | 3 | 4 |
| Siparunaceae | *Siparuna guianensis* | Plot-weighted | Small | | 1.0 | 3 | 4 |
| Euphorbiaceae | *Sagotia racemosa* | Plot-weighted | Small | 1.0 | 1.0 | 2 | 1 |
| Lecythidaceae | *Eschweilera coriacea* | Plot-weighted | Small | 1.5 | 1.0 | 3 | 4 |
| Lecythidaceae | *Couratari stellata* | Plot-weighted | Small | | 1.0 | 3 | 2 |
| Solanaceae | *Solanum crinitum* | Plot-weighted | Small | 0.8 | 1.0 | 1 | 2 |
| Lecythidaceae | *Gustavia augusta* | Plot-weighted | Small | | 1.0 | 2 | 3 |
| Lecythidaceae | *Lecythis lurida* | Plot-weighted | Small | 1.3 | 1.0 | 3 | 3 |

| | | | | | | | |
|---|---|---|---|---|---|---|---|
| Fabaceae | *Swartzia laurifolia* | Plot-weighted | Small | 1.3 | 0.9 | 3 | 4 |
| Boraginaceae | *Cordia exaltata* | Plot-weighted | Small | | 0.8 | 3 | 4 |
| Euphorbiaceae | *Croton matourensis* | Plot-weighted | Small | 0.9 | 0.7 | 2 | 3 |
| Euphorbiaceae | *Mabea angustifolia* | Plot-weighted | Small | | 0.7 | 3 | 4 |
| Achariaceae | *Lindackeria paraensis* | Plot-weighted | Small | | 0.7 | 3 | 4 |
| Apocynaceae | *Ambelania acida* | Plot-weighted | Small | | 0.7 | 3 | 3 |
| Myrtaceae | *Myrciaria floribunda* | Plot-weighted | Small | 0.8 | 0.7 | 2 | 3 |
| Lacistemataceae | *Lacistema aggregatum* | Plot-weighted | Small | | 0.7 | 3 | 3 |
| Melastomataceae | *Miconia minutiflora* | Plot-weighted | Small | | 0.7 | 2 | 3 |
| Anacardiaceae | *Tapirira guianensis* | Plot-weighted | Small | 0.8 | | 3 | 4 |
| Myrtaceae | *Myrcia cuprea* | Plot-weighted | Small | 1.0 | | 2 | 2 |
| Sapotaceae | *Pouteria macrophylla* | Plot-weighted | Small | 0.8 | | 3 | 3 |
| Salicaceae | *Casearia decandra* | Plot-weighted | Small | 0.8 | | 3 | 4 |
| Phyllanthaceae | *Margaritaria nobilis* | Plot-weighted | Small | 0.7 | | 2 | 3 |
| Fabaceae | *Ormosia paraensis* | Plot-weighted | Small | 1.0 | | 3 | 3 |

| | | | | | | | |
|---|---|---|---|---|---|---|---|
| Fabaceae | *Inga capitata* | Plot-weighted | Small | 0.8 | | 3 | 4 |
| Myrtaceae | *Marlierea spruceana* | Plot-weighted | Small | 0.8 | | 1 | 1 |
| Fabaceae | *Inga thibaudiana* | Plot-weighted | Small | 0.8 | | 3 | 4 |
| Euphorbiaceae | *Maprounea guianensis* | Plot-weighted | Small | 0.9 | | 3 | 4 |
| Fabaceae | *Bauhinia acreana* | Plot-weighted | Small | 0.8 | | 3 | 2 |

### References


Fauset, S., Johnson, M. O., Gloor, M., Baker, T. R., Monteagudo M., A., Brienen, R. J. W., Feldpausch, T. R., Lopez-Gonzalez, G., Malhi, Y., ter Steege, H., Pitman, N. C. A., Baraloto, C., Engel, J., Pétronelli, P., Andrade, A., Camargo, J. L. C., Laurance, S. G. W., Laurance, W. F., Chave, J., … Phillips, O. L. (2015). Hyperdominance in Amazonian forest carbon cycling. *Nature Communications*, *6*(1), Article 1. https://doi.org/10.1038/ncomms7857

Funk, C., Peterson, P., Landsfeld, M., Pedreros, D., Verdin, J., Shukla, S., Husak, G., Rowland, J., Harrison, L., Hoell, A., & Michaelsen, J. (2015). The climate hazards infrared precipitation with stations—A new environmental record for monitoring extremes. *Scientific Data*, *2*(1), Article 1. https://doi.org/10.1038/sdata.2015.66

MapBiomas, M. (2024). *Collection 9 of the Annual Land Cover and Land Use Maps of Brazil (1985-2023)* [Dataset]. MapBiomas Data. https://doi.org/10.58053/MapBiomas/XXUKA8

ter Steege, H., Pitman, N. C. A., Sabatier, D., Baraloto, C., Salomão, R. P., Guevara, J. E., Phillips, O. L., Castilho, C. V., Magnusson, W. E., Molino, J.-F., Monteagudo, A., Núñez Vargas, P., Montero, J. C., Feldpausch, T. R., Coronado, E. N. H., Killeen, T. J., Mostacedo, B., Vasquez, R., Assis, R. L., … Silman, M. R. (2013). Hyperdominance in the Amazonian Tree Flora. *Science*, *342*(6156), 1243092. https://doi.org/10.1126/science.1243092
